# Macropinocytosis-Mediated Recyclable LYTACs (McR-TACs) for Membrane and Extracellular Protein Degradation

**DOI:** 10.1101/2025.09.08.674971

**Authors:** Peixin Liu, Yule Li, Yawen You, Yu Chen, Michelle Cai, Quanyin Hu

## Abstract

The degradation of cell membrane and extracellular proteins with lysosome-targeting chimeras (LYTACs) is hampered by non-recyclable and lysosome-shuttling receptor-dependent mechanisms, limiting the wide application of this emerging technology. Here, we developed macropinocytosis-mediated recyclable LYTACs (McR-TACs) that could durably degrade cell membrane and extracellular proteins in a receptor-independent manner. This McR-TAC platform demonstrated the feasibility of harnessing endogenous transport pathways for developing next-generation recyclable protein degraders with broad applicability.

---

Lysosome-targeting chimaeras (LYTACs) have emerged as a transformative modality for the targeted degradation of diverse therapeutic targets, including membrane and extracellular proteins^1-3^. However, a critical limitation of the current LYTAC platform lies in its concomitant degradation alongside the protein of interest (POI) during lysosomal trafficking^4^. In contrast to the catalytic degradation mechanism of proteolytic targeting chimeras, LYTACs usually demonstrate very limited recyclability in targeted protein degradation (TPD)^5^. Since the initial development of LYTACs, their recyclability lacks systematic and intensive exploration. In the current TPD field, there have been proposed eager demands for recyclable LYTACs, owing to their theoretical advantages of avoiding frequent administration while maintaining the potency^6, 7^. The most recent advancements are only achieved by indirect mechanisms with more focus on recycling the endocytic receptors rather than LYTAC itself, such as designing degraders using the pH-sensitive mechanism of recycling neonatal Fc receptor^8-10^. Similar to conventional LYTACs, this strategy predominantly depends on the engagement of lysosomal shuttle receptors for endocytosis. However, due to the potential competition from endogenous ligands (e.g., mannose-6-phosphate-modified glycoproteins), the in vivo efficiency of LYTACs might be affected due to individual differences when applied on a large scale^11^. Besides, the targeting of endocytic receptors is complicated by their heterogeneous expression across various tissues and cell types^12^. The number of highly specific receptors, such as the exclusive expression of ASGPR in hepatocytes, is very limited^7^. Moreover, the intricate endosomal microenvironment and the ambiguous sorting mechanisms of endocytic cargo complicate the design of recyclable LYTACs^13^. These limitations place even higher demands on the structural design of recyclable LYTACs, which will play a significant role in promoting the extensive development and application of LYTACs in the future. Consequently, the development of next-generation LYTACs platforms to overcome the constraints of non-recyclable and receptor-dependent principles is of great significance in the TPD field.

Mechanistic decoding of recyclable LYTACs hinges on system-level investigation of cellular trafficking networks. Cells possess an intrinsic recycling mechanism known as transcytosis, allowing substrates to be transported intactly^14^. Transcytosis employs multiple endocytic pathways, among which macropinocytosis represents a unique receptor-independent cellular uptake mechanism^15^. Characterized by actin-driven membrane ruffles, micropinocytosis mediates the internalization of solute molecules, nutrients and antigens. For example, albumin can be extensively internalized by tumor cells through macropinocytosis to sustain their high-level nutritional demands, particularly in KRAS-mutated pancreatic cancers^16^. This endogenous intracellular transport pathway inspires us to envision the design of new LYTACs that can trigger macropinocytosis, enabling the recyclability of molecules without relying on recycling receptors. This strategy may promote the development of next-generation recyclable and receptor-independent LYTACs, with potential applications in various disease therapies.

To achieve this goal, we proposed a novel platform design for macropinocytosis-mediated recyclable LYTACs (McR-TACs). Specifically, McR-TACs were designed as polymer-drug conjugates where POI ligands were covalently attached to a polymer capable of triggering macropinocytosis. This polymer, synthesized through reversible addition-fragmentation chain transfer (RAFT) polymerization of zwitterionic tertiary amine oxides, has been demonstrated to interact with the hydrophilic head groups of phospholipids on cell membranes, thereby inducing internalization (**Supplementary Fig. 1-5**)^17, 18^. To demonstrate the TPD capability of McR-TACs, we chose the programmed cell death-ligand 1 (PD-L1) as a targeted membrane protein for degradation verification. PD-L1 commonly expresses on the surface of tumor cells, facilitating immune evasion and serving as a prevalent target in clinical cancer immunotherapy^19^. We used BMS-1166, a small molecule inhibitor of PD-L1, as the ligand to develop McR-TACs_PD-L1_ (**Fig. 1a**). Considering that the length of LYTACs molecules has an important influence on their protein degradation efficiency, we synthesized and screened McR-TACs_PD-L1_ analogs of varying lengths. The length was controlled by adjusting the ratio of tertiary amine oxide monomers (x=∼10, 20, 40, 80 and 120) in the RAFT polymerization while keeping the amount of POI ligands constant (y=∼5). We treated human breast cancer cells MDA-MB-231 with McR-TACs_PD-L1_ analogs of different lengths and concentrations, and collected cell lysates for immunoblot analysis after 24 h. These results indicated that as the polymer length of McR-TACs_PD-L1_ increased, PD-L1 degradation was observed when the number of macropinocytosis-triggering ligands reached approximately 40 (**Fig. 1b-c**). Further increasing the monomer number to around 80 yielded optimal degradation efficiencies among all screened McR-TACs_PD-L1_. Additionally, McR-TACs_PD-L1_ (x=∼80, y=∼5) also induced a concentration- and time-dependent downregulation of PD-L1 expression (**Fig. 1d** and **Supplementary Fig. 6**). Thus, we used the optimized McR-TACs (x=∼80, y=∼5) for the following studies. To further confirm that PD-L1 degradation occurs through the lysosomal pathway, we investigated the localization of PD-L1 in MDA-MB-231 cells treated with McR-TACs_PD-L1_ for 8 h. Fluorescent labeling of the McR-TACs_PD-L1_ was characterized by fluorescent spectrum scanning and did not significantly affect its protein degradation efficiency (**Supplementary Fig. 7**). Compared to the untreated group, where PD-L1 signals were predominantly located on the cell membrane, McR-TACs_PD-L1_ treatment resulted in PD-L1 signals migrating from the cell membrane into the cytoplasm and co-localized with lysosomes marked by the lysosomal marker LAMP1 (**Fig. 1e** and **Supplementary Fig. 8**). These results demonstrated that McR-TACs_PD-L1_ could effectively mediate the lysosomal degradation of membrane protein PD-L1.

**Fig. 1:**
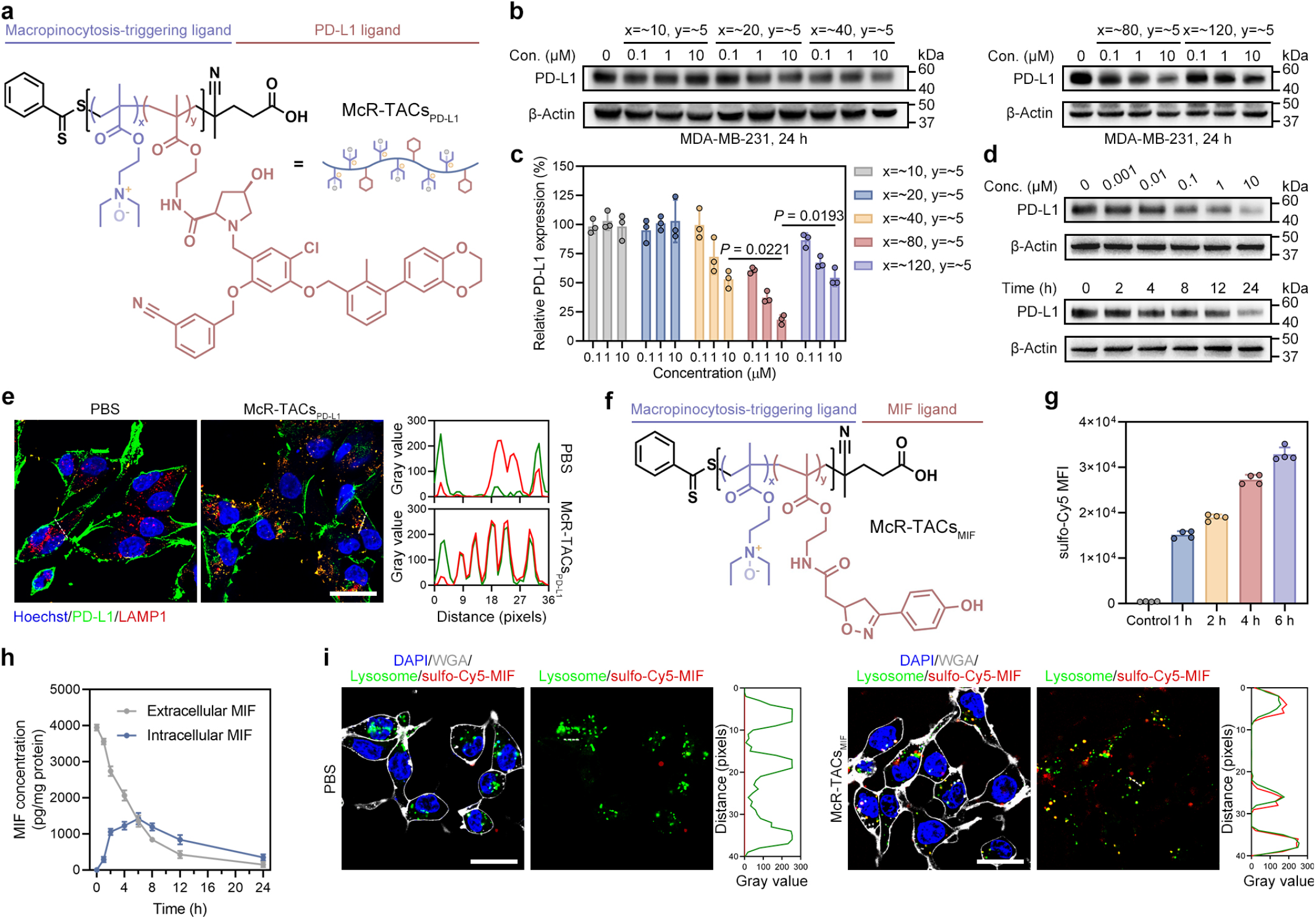
McR-TACs transport membrane and extracellular proteins to lysosomes for degradation. **a**, Chemical structure of McR-TACs for PD-L1 degradation. **b**, Western blot analysis of PD-L1 degradation in MDA-MB-231 cells treated with different lengths of McR-TACs_PD-L1_ under indicated concentrations for 24 h. **c**, Semi-quantification of relative PD-L1 levels in **b** (*n* = 3 per group). Data are presented as the mean values ± s.d. and were analyzed by a two-way ANOVA, followed by a Tukey’s multiple comparisons test. **d**, Western blot analysis of PD-L1 degradation in MDA-MB-231 cells treated with McR-TACs_PD-L1_ under indicated concentrations or times. **e**, Confocal imaging of the co-localization between PD-L1 and the lysosome marker LAMP1 in MDA-MB-231 cells upon 8 h treatment with PBS or 10 μM of McR-TACs_PD-L1_. Scale bar, 40 μm. Plot profiles of the overlapped LAMP1 and PD-L1 signals are indicated by dotted white lines in confocal images. **f**, Chemical structure of McR-TACs for MIF degradation. **g**, Flow cytometry analysis of sulfo-Cy5-MIF internalization in MDA-MB-231 cells at indicated times (*n* = 4 per group). Data are presented as the mean values ± s.d. **h**, The degradation kinetics of extracellular and intracellular MIF in 4T1 cells quantified with ELISA (*n* = 3 per group). Data are presented as the mean values ± s.d. **i**, Confocal imaging of the co-localization between sulfo-Cy5-MIF and lysosomes in MDA-MB-231 cells upon 4 h treatment with PBS or 10 μM McR-TACs_MIF_. Scale bar, 40 μm. Plot profiles of the overlapped lysosome and sulfo-Cy5-MIF signals are indicated by dotted white lines in confocal images.

To further investigate whether McR-TACs can be applied for extracellular protein degradation, we incorporated the macrophage migration inhibitory factor (MIF) inhibitor ISO-1 as warhead to develop McR-TACs_MIF_ (**Fig. 1f** and **Supplementary Fig. 9-12**). MIF, a cytokine typically released under inflammatory conditions, binds to immune cells to suppress immune responses^20^. This extracellular protein has been utilized as a POI in the design of LYTACs^21^. We added sulfo-Cy5-labeled human MIF with McR-TACs_MIF_ to the culture medium of MDA-MB-231 cells. The uptake of MIF driven by McR-TACs_MIF_ was characterized by flow cytometry (**Fig. 1g**). In order to quantify the MIF degradation in vitro and avoid the interference of endogenous MIF, McR-TACs_MIF_ and MIF were added to the medium of murine 4T1 cells. Both intracellular and extracellular concentration of MIF were quantified by ELISA. Results showed that MIF could be quickly internalized with McR-TACs_MIF_ for subsequent degradation (**Fig. 1h**). Additionally, we also observed that the internalized MIF signals co-localized with lysosomes (**Fig. 1i**). Since the warhead alone treatment group (BMS-1166 or ISO-1) could not induce the POI degradation (**Supplementary Fig. 13**), we demonstrated that McR-TACs showed promising potency for the degradation of both membrane and extracellular proteins.

Subsequently, we tried to figure out whether the intracellular trafficking of McR-TACs worked through the macropinocytosis pathway as hypothesized. First, the POI ligands were replaced with Cy5 to investigate the transportation of polymers independent of warheads. The intracellular location of polymers was monitored in living MDA-MB-231 cells with organelle stainings (**Extended Data Fig. 1**). Confocal images showed that polymers were first transported into macropinosomes and then escaped from them during 1 to 4 hours. Afterwards, polymers showed co-localization with endoplasmic reticulum (ER) and moved to Golgi apparatus from 4 to 8 hours. Throughout the entire observation period, polymers did not colocalize with lysosomes. Pearson’s R values were used to indicate the dynamic changes in the co-localization of polymers with organelles. Based on the above pathway, we began by investigating the cellular uptake mechanism of McR-TACs. Cells were pretreated with various inhibitory conditions for endocytosis, including caveolae-mediated endocytosis inhibitor (filipin), clathrin-mediated endocytosis inhibitor (chlorpromazine), macropinocytosis-mediated endocytosis inhibitor (EIPA), free warheads (BMS-1166), micropinocytosis-related protein knockdown (RAC1 siRNA) and low temperature (4°C), followed by co-incubation with sulfo-Cy5-McR-TACs_PD-L1_^22^. Flow cytometry results revealed that the uptake of sulfo-Cy5-McR-TACs_PD-L1_ by MDA-MB-231 cells primarily was through the macropinocytosis pathway, indicated by the attenuated uptake after EIPA and RAC1 siRNA pre-treatment (**Fig. 2a** and **Supplementary Fig. 14**). Next, we examined the intracellular transportation of McR-TACs_PD-L1_. After 2 h of incubation, McR-TACs_PD-L1_ exhibited significant co-localization with macropinosomes rather than lysosomes (**Fig. 2b** and **Supplementary Fig. 15**). This observation provides critical evidence for the recyclability of McR-TACs_PD-L1_, as most degrader signals did not directly enter lysosomes along with the POI which is a typical pathway for traditional LYTACs. Upon extending the incubation time to 4 and 8 hours, the fluorescence signals of McR-TACs_PD-L1_ further co-localized with the ER and Golgi apparatus, which are known to be the organelles involved in processing secretory cargo^23^. This suggests that McR-TACs_PD-L1_ may be excreted by cells via exocytosis, enabling its recyclable degradation capability. Similar uptake mechanism and intracellular pathway were also observed in MIF degradation by McR-TACs_MIF_ (**Fig. 2c** and **Supplementary Fig. 16-17**). To further illustrate the impact of each intracellular trafficking step of McR-TACs on TPD, we inhibited specific transport processes, including POI binding, endocytosis, lysosomal degradation, and exocytosis. Western blot analysis demonstrated that the combination of McR-TACs_PD-L1_ with BMS-1166 (POI ligand), EIPA, chloroquine (CQ, lysosomal inhibitor), or Exo1 (exocytosis inhibitor) resulted in attenuated PD-L1 degradation efficiency in MDA-MB-231 cells (**Fig. 2d**). A similar phenomenon was also observed in MIF degradation (**Supplementary Fig. 18**). These findings demonstrated that McR-TACs mediated intracellular trafficking and lysosomal degradation of POI through the macropinocytosis pathway. In order to investigate the McR-TACs’ performance on different cell lines, we employed four more cell lines in addition to MDA-MB-231 and 4T1-hPD-L1 cells used in this study with different macropinocytosis levels which are positively correlated with the RAS expression. U87 and PANC-1 cells showed high expression of RAS while BxPC-3 and Caco-2 cells showed low expression of RAS (**Extended Data Fig. 2**). The uptake of McR-TACs_PD-L1_ on these cells positively correlated with RAS expression levels. Pre-treatment with KRAS siRNA could compromise the uptake of McR-TACs_PD-L1_ by the cells (U87, 4T1-hPD-L1, MDA-MB-231 and PANC-1) with high RAS expression.

**Fig. 2:**
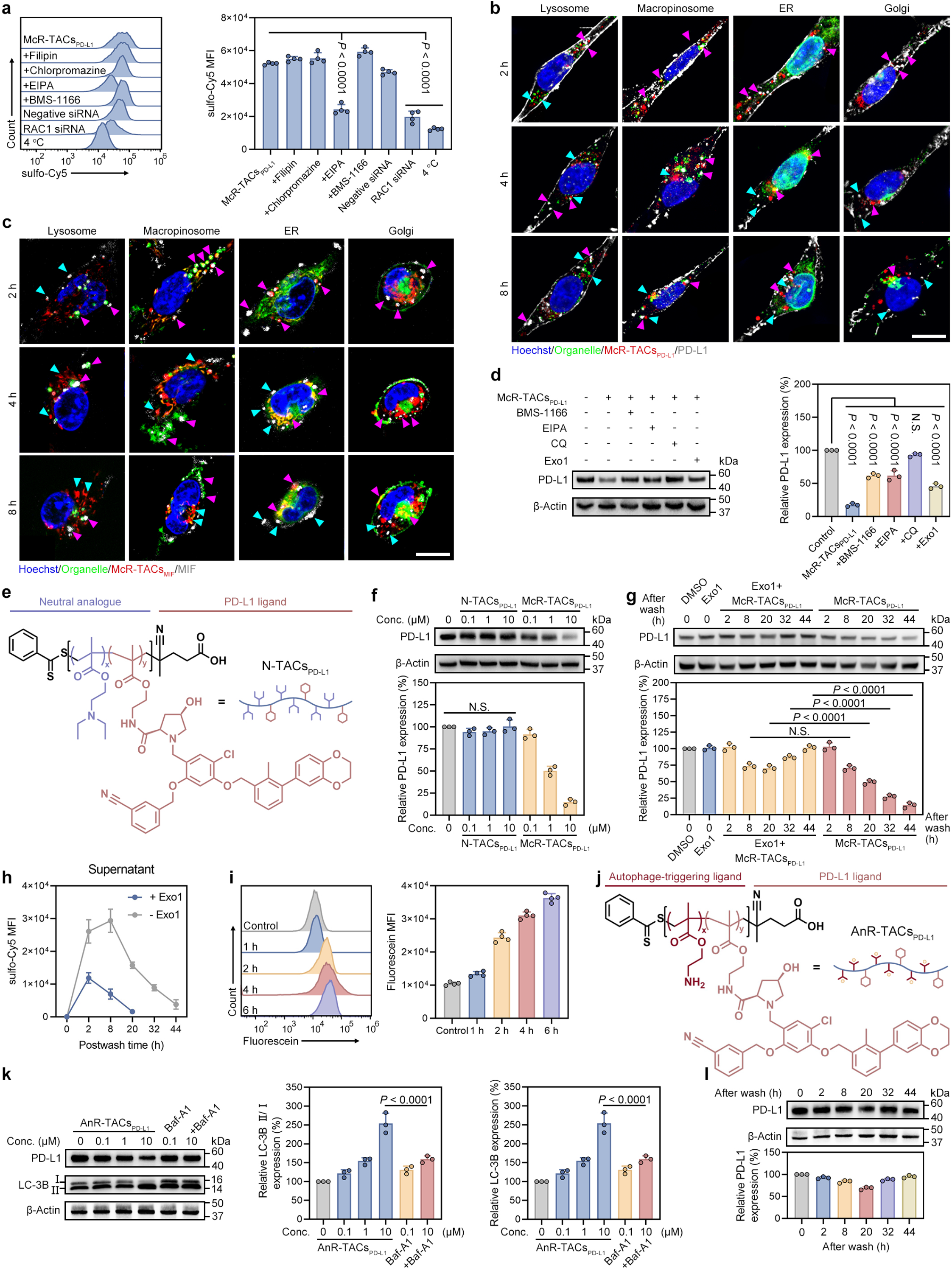
McR-TACs show zwitterionic structure-dependent and recyclable protein degradation effects through the macropinocytosis pathway. **a**, Flow cytometry analysis of sulfo-Cy5-McR-TACs_PD-L1_ internalization in MDA-MB-231 cells pre-treated with indicated uptake inhibitors (*n* = 4 per group). **b**, Confocal imaging of the localization of sulfo-Cy5-McR-TACs_PD-L1_, PD-L1 and organelles including lysosomes stained with anti-LAMP1, macropinosomes labeled with Fluorescein-BSA, ER stained with anti-Calnexin and Golgi stained with anti-GM130 in MDA-MB-231 cells at indicated times. Scale bar, 20 μm. **c**, Confocal imaging of the localization of sulfo-Cy5-McR-TACs_MIF_, Fluorescein-MIF and organelles including lysosomes labeled with lysotracker red, macropinosomes labeled with TRITC-dextran, ER labeled with ER tracker red and Golgi labeled with TMR ceramide in MDA-MB-231 cells at indicated times. Scale bar, 20 μm. **d**, Western blot and semi-quantified analysis of PD-L1 degradation in MDA-MB-231 cells treated with McR-TACs_PD-L1_ or McR-TACs_PD-L1_ plus BMS-1166, EIPA, CQ or Exo1 at 10 μM for 24 h (*n* = 3 per group). **e**, Chemical structure of N-TACs_PD-L1_. **f**, Western blot and semi-quantified analysis of PD-L1 degradation in MDA-MB-231 cells treated with N-TACs_PD-L1_ or McR-TACs_PD-L1_ under indicated concentrations for 24 h (*n* = 3 per group). **g**, Western blot and semi-quantified analysis of recyclable PD-L1 degradation in MDA-MB-231 cells pre-treated with 20 μM McR-TACs_PD-L1_ with or without Exo1 for 4 h and further incubated in fresh medium with or without Exo1 for indicated time points (*n* = 3 per group). **h**, Quantified sulfo-Cy5-McR-TACs_PD-L1_ fluorescent intensity in the supernatant of MDA-MB-231 cells at indicated time points after washout (*n* = 3 per group). Data are presented as the mean values ± s.d. **i**, Flow cytometry analysis of Fluorescein-MIF internalization in MDA-MB-231 cells after washing out McR-TACs_MIF_ and sulfo-Cy5-MIF (*n* = 4 per group). Data are presented as the mean values ± s.d. **j**, Chemical structure of AnR-TACs for PD-L1 degradation. **k**, Western blot and semi-quantified analysis of PD-L1 degradation and autophagy levels in MDA-MB-231 cells treated with AnR-TACs_PD-L1_ or AnR-TACs_PD-L1_ plus Baf-A1 under indicated concentrations (*n* = 3 per group). **l**, Western blot and semi-quantified analysis of PD-L1 degradation in MDA-MB-231 cells pre-treated with 20 μM AnR-TACs_PD-L1_ for 4 h and further incubated in fresh medium for indicated times (*n* = 3 per group). Data are presented as the mean values ± s.d. For **a** and **d**, data are presented as the mean values ± s.d. and were analyzed by a one-way ANOVA, followed by a Tukey’s multiple comparisons test. For **f, g** and **k**, data are presented as the mean values ± s.d. and were analyzed by a two-way ANOVA, followed by a Tukey’s multiple comparisons test.

Since the zwitterionic structure of McR-TACs could interact with cell membrane, we hypothesized that the tertiary amine oxide of McR-TACs is crucial for triggering macropinocytosis and subsequent protein degradation. To prove this, we synthesized a neutral analog of McR-TACs_PD-L1_ (N-TACs_PD-L1_), which differs primarily in the tertiary amine without oxidation (**Fig. 2e** and **Supplementary Fig. 19-21**). In contrast to McR-TACs_PD-L1_, N-TACs_PD-L1_ failed to induce significant PD-L1 degradation under various concentrations, indicating that the TPD of McR-TACs is zwitterion-dependent (**Fig. 2f**).

After exploring the biological and chemical mechanisms of McR-TACs, we further investigated whether McR-TACs can be recycled for protein degradation. We first conducted washout experiments to test the duration of PD-L1 degradation. MDA-MB-231 cells were pretreated with 20 μM McR-TACs_PD-L1_ for 4 h, after which the degraders-containing medium was replaced with fresh medium, and the PD-L1 expression was measured at different time points. The fluorescent intensity of sulfo-Cy5-McR-TACs_PD-L1_ in both cell and supernatant were also monitored. Exo1 was used as a control to impair recycling exocytosis. McR-TACs_PD-L1_ demonstrated sustained PD-L1 degradation efficacy. Even after 44 h of washout, McR-TACs_PD-L1_ were still capable of degrading approximately 90% of PD-L1 (**Fig. 2g**). However, Exo1 disrupted this recyclable degradation process, attenuating the degradation efficiency within 24 h and allowing PD-L1 expression to gradually recover. In addition, the results of McR-TACs_PD-L1_ kinetics revealed that McR-TACs_PD-L1_ could be effectively excreted from cells and maintained in the system for an extended period (**Fig. 2h**). However, this recycling process was inhibited by Exo1, which could not affect the internalization of McR-TACs but resulted in the entrapment and subsequent degradation of McR-TACs_PD-L1_ within cells (**Supplementary Fig. 22-23**). Then, we attempted to investigate the feasibility of recyclable MIF degradation. McR-TACs_MIF_ and sulfo-Cy5-MIF were mixed to treat MDA-MB-231 cells for 4 hours and followed by being washed out. Fluorescein-MIF was added to the fresh medium without McR-TACs_MIF_ for further incubation. The intensity of both internalized sulfo-Cy5-MIF and newly added Fluorescein-MIF were monitored by flow cytometry (**Fig. 2i** and **Supplementary Fig. 24**). The decreased intensity of sulfo-Cy5-MIF and increased intensity of Fluorescein-MIF indicated the recyclable degradation of MIF by McR-TACs_MIF_. These findings collectively demonstrate that McR-TACs possess recyclable properties for TPD.

Given that Exo1 did not completely inhibit the exocytosis process of McR-TACs, we sought to further demonstrate the recyclable degradation efficiency by designing a McR-TACs_PD-L1_ analog that could degrade POI through a non-recyclable pathway. Recent study reported that cationic AUTABs could degrade POI through the autophagy pathway^24^. For receptor-independent LYTACs, their charged property plays a significant role in the endocytic pathway. Therefore, we synthesized a cationic autophagy-mediated non-recyclable LYTACs (AnR-TACs_PD-L1_) (**Fig. 2j**). The major modification of degrader structure was to replace the tertiary amine oxide monomer with a primary amine monomer (**Supplementary Fig. 25-28**). Western blot analysis showed that AnR-TACs_PD-L1_ exhibited concentration-dependent PD-L1 degradation, which could be attenuated by the autophagolysosome inhibitor Bafilomycin A1 (Baf-A1) (**Fig. 2k**). Additionally, we observed that AnR-TACs_PD-L1_ upregulated the expression of LC-3B, a marker of autophagy activation. Imaging of intracellular autophagosomes further confirmed the ability of AnR-TACs_PD-L1_ to activate autophagy (**Supplementary Fig. 29**). In contrast, no autophagy activation was observed in other treatment groups with LYTACs. Having demonstrated that AnR-TACs_PD-L1_ degraded PD-L1 via the autophagy pathway, we conducted the same washout experiment to compare the durability of its degradation effects. Western blot results showed that AnR-TACs_PD-L1_ maintained only approximately 30% PD-L1 degradation efficacy within the first 24 hours, after which PD-L1 expression gradually recovered (**Fig. 2l**). By comparison, we concluded that AnR-TACs_PD-L1_, due to its non-recyclable autophagy-mediated degradation mechanism, could not sustain effective degradation of membrane proteins after washout. In contrast, McR-TACs showed recyclable degradation through the macropinocytosis pathway (**Extended Data Fig. 4**).

The number-averaged molecular weight (Mn) and polydispersity index (PDI) of each polymer was characterized by gel permeation chromatography (GPC) (**Supplementary Fig. 30** and **Table. 1**). In addition, the number of two monomers (x and y) in the polymer structure could be estimated according to the ^1^H-NMR spectrum and GPC results (**Supplementary Table. 2**). The in vitro cytotoxicity of warheads and different groups of LYTACs was investigated in MDA-MB-231 cells (**Supplementary Fig. 31**). McR-TACs and N-TACs showed good biocompatibility within the concentration range used in this study. While AnR-TACs exhibited potential cytotoxicity at high concentrations due to their positive charge.

Encouraged by the recyclable degradation efficiency of McR-TACs in vitro, we continued to test their in vivo TPD efficiency. When applied to the in vivo conditions, the influence of extracellular context on the behavior of degraders needs to be studied first. Considering that the acidic tumor microenvironment may have an impact on the behavior of charge-dependent degraders, the PD-L1 degradation of McR-TACs_PD-L1_ under different pH conditions (physiological pH 7.4 and tumor microenvironment pH 6.8) was investigated (**Extended Data Fig. 3a-b**). There are no significant changes in the protein degradation efficiency between two different pH conditions. Besides the impact of pH, we also evaluated the impact of the complex components in the blood circulation, including blood cells and serum proteins, on the performance of McR-TACs after systemic injection. Red blood cells were incubated with sulfo-Cy5-McR-TACs_PD-L1_ and then detected under confocal microscopy and flow cytometry (**Extended Data Fig. 3c-e**). Results demonstrated that McR-TACs_PD-L1_ could bind to the red blood cells and be washed away by repeated syringe extraction that mimics the shear force in the bloodstream. This reversible adsorption allowed the McR-TACs’ transportation from red blood cells to tumor tissue, a mechanism reported before for the zwitterionic polymers^25^. Further, McR-TACs_PD-L1_ did not show obvious stickiness to the serum protein with BSA as a model serum protein (**Extended Data Fig. 3f-h**).

Since PD-L1 degradation disrupts the PD-1/PD-L1 interaction between cancer cells and immune cells to enhance antitumor immunity, we used a luciferase-transfected 4T1 murine breast cancer cell line with human PD-L1 expression (4T1-hPD-L1 cells) to establish a triple-negative breast cancer (TNBC) mouse model. We first investigated the in vivo kinetics of McR-TACs_PD-L1_ and AnR-TACs_PD-L1_ after the i.t. injection. The dynamic fluorescence signals of free probes and fluorescently labeled degraders were monitored via in vivo imaging system (IVIS) (**Fig. 3a**). IVIS results demonstrated rapid clearance of free probes due to unrestricted diffusion. In contrast, both AnR-TACs_PD-L1_ and McR-TACs_PD-L1_ exhibited extended tumor retention. McR-TACs_PD-L1_ showed significantly prolonged i.t. accumulation compared to AnR-TACs_PD-L1_ (**Fig. 3b**). Co-injection of exocytosis inhibitor Exo1 markedly reduced McR-TACs_PD-L1_ accumulation in tumors, confirming the recycling function.

**Fig. 3:**
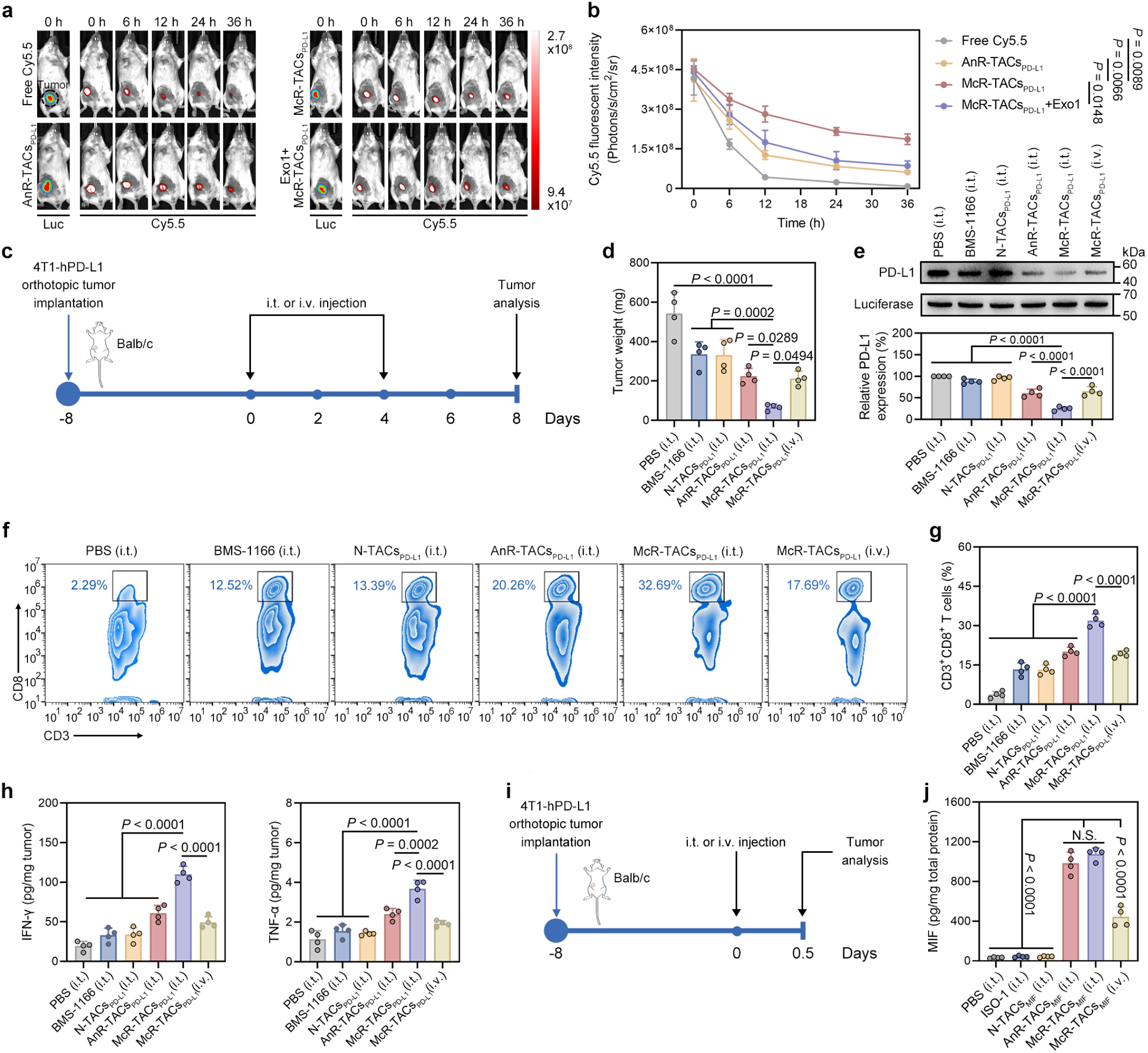
McR-TACs degrade targeted proteins and enhance antitumor immune responses in an orthotopic TNBC mouse model. **a**, IVIS imaging of tumor-bearing mice at indicated time points after i.t. injection of Cy5.5, sulfo-Cy5.5-labeled AnR-TACs_PD-L1_, McR-TACs_PD-L1_ or McR-TACs_PD-L1_ plus Exo1 at a dose of 0.2 mg/kg sulfo-Cy5.5 (*n* = 4 mice). **b**, Quantified Cy5.5 fluorescent intensity of tumor sites at indicated time points (*n* = 4 mice). Data are presented as the mean values ± s.e.m. and were analyzed by a two-way ANOVA, followed by a Tukey’s multiple comparisons test. **c**, Schematic illustration of the treatment plan in the TNBC mice model with i.t. injection of PBS, BMS-1166, N-TACs_PD-L1_, AnR-TACs_PD-L1_, McR-TACs_PD-L1_ or i.v. injection of McR-TACs_PD-L1_ at a dose of 3 mg/kg BMS-1166 (*n* = 8 mice). **d**, The tumor weight measurements on day 8 after treatments (*n* = 4 mice). **e**, Western blot and semi-quantified analysis of PD-L1 degradation in the tumor tissues on day 8 after treatments (*n* = 4 mice). **f**, Flow cytometry analysis of CD8^+^ T cell proportions out of the total CD45^+^ CD3^+^ cell population (*n* = 4 mice). **g**, Proportions of CD8^+^ T cells out of the total CD45^+^ CD3^+^ cell population in the tumor tissues on day 8 after treatments (*n* = 4 mice). **h**, Quantification of IFN-γ and TNF-α levels in the tumor tissues by ELISA on day 8 after treatments (*n* = 4 mice). **i**, Schematic illustration of the treatments in the TNBC mice model with i.t. injection of MIF plus PBS, ISO-1, N-TACs_MIF_, AnR-TACs_MIF_, McR-TACs_MIF_ or i.v. injection of McR-TACs_MIF_ at a dose of 0.2 mg/kg ISO-1 (*n* = 4 mice). **j**, Quantification of intracellular MIF levels of the tumor tissues by ELISA 12 h after injection (*n* = 4 mice). For **d, e, g, h** and **j**, data are presented as the mean values ± s.d. and were analyzed by a one-way ANOVA, followed by a Tukey’s multiple comparisons test.

Subsequently, we evaluated the therapeutic efficacy of McR-TACs_PD-L1_ in tumor suppression and immune activation. Due to the potential cytotoxicity of AcR-TACs and poor solubility of N-TACs, the efficacy of these two degraders was only studied by intratumoral (i.t.) injection. While McR-TACs were administered through both i.t. and i.v. routes. When tumor volumes reached approximately 100 mm^3^, mice received i.t. or i.v. injections (once every four days for a total of two times) of PBS, BMS-1166, N-TACs_PD-L1_, AnR-TACs_PD-L1_, or McR-TACs_PD-L1_ (**Fig. 3c**). By day 8 post-treatment, McR-TACs_PD-L1_ showed potent tumor suppression indicated by the tumor weights (**Fig. 3d** and **Supplementary Fig. 32**). Western blot analysis revealed McR-TACs_PD-L1_ through i.t. injection achieved the most potent in vivo PD-L1 degradation efficiency (approximately 70%) (**Fig. 3e**). Flow cytometric profiling of tumor-infiltrating lymphocytes showed that McR-TACs_PD-L1_-treated tumors exhibited a higher proportion of CD45^+^ CD3^+^ CD8^+^ T cells compared to other treatment groups (**Fig. 3f-g** and **Supplementary Fig. 33**). In addition, elevated levels of immunostimulatory cytokines, including interferon-γ (IFN-γ) and tumor necrosis factor-α (TNF-α), were detected in McR-TACs_PD-L1_-treated tumors, indicating enhanced antitumor immune activation (**Fig. 3h**).

Notably, no significant body weight changes were observed in any treatment group during the treatment (**Supplementary Fig. 34**). Hematological analysis conducted one-week post-treatment revealed comparable blood parameters across all groups, confirming the favorable safety profile of McR-TACs_PD-L1_ (**Supplementary Fig. 35**). These findings collectively demonstrated that McR-TACs_PD-L1_ effectively degraded PD-L1 in vivo, subsequently activated antitumor immunity through T cell infiltration and cytokine induction while maintaining good safety profile.

To further validate the extracellular protein degradation capability of McR-TACs_MIF_ in vivo, we explored its feasibility using human MIF as the POI. To avoid interference from endogenous murine MIF in the tumor microenvironment, human MIF was administered through i.t. or i.v. injections with different LYTACs (**Fig. 3i**). The synthesis of N-TACs_MIF_ and AnR-TACs_MIF_ control groups for MIF degradation was similar to that of McR-TACs_MIF_ (**Supplementary Fig. 36-39**). Given the rapid clearance of human MIF in mice, degradation assessment was limited to short-term analysis. Tumors were harvested 12 hours post-injection, and single-cell suspensions were prepared for intracellular human MIF quantification via ELISA. Both AnR-TACs_MIF_ and McR-TACs_MIF_ significantly enhanced human MIF internalization compared to PBS, ISO-1 and N-TACs_MIF_ treatment groups, verifying the protein degradation capability of extracellular proteins with McR-TACs_MIF_ in vivo (**Fig. 3j**).

In summary, we developed a recyclable LYTACs platform (McR-TACs) inspired by the endogenous macropinocytosis pathway. McR-TACs enabled receptor-independent degradation of membrane and extracellular proteins, demonstrating promising and durable degradation efficiency both in vitro and in vivo. Notably, McR-TACs enhanced anti-tumor immunity with a favorable safety profile in a TNBC mouse model. Collectively, McR-TACs addressed critical limitations of conventional LYTACs, such as non-recyclable degraders and receptor-dependent endocytosis, while offering a strategic blueprint for the next-generation LYTAC development.

## Methods

### Materials

All the chemicals for synthesis were purchased from AmBeed unless otherwise listed. All the anhydrous solvents for synthesis and siRNA (negative, RAC1, KRAS siRNA) were purchased from Thermo Fisher. Cell uptake, transportation and exocytosis inhibitors and human MIF protein were purchased from Cayman Chemical. All the fluorescent probes were purchased from Lumiprobe Corporation unless otherwise listed. Primary and secondary antibodies for western blot and confocal imaging were obtained from Cell Signaling Technology or Proteintech. Primary antibodies for flow cytometry and ELISA kits were obtained from BioLegend.

### Cell lines and animals

The human MDA-MB-231 breast cancer cell line and murine 4T1 breast cancer cell line were obtained from ATCC. The 4T1-hPD-L1 cell line was gifted by S.-O. Lim (Purdue University). Cells were cultured in a CO_2_ incubator (Fisher) at 37 °C with 5% CO_2_ and 90% relative humidity. Six-week-old female Balb/c mice were purchased from the Jackson Laboratory and were housed in a pathogen-free facility under a 12-h light/12-h dark cycle at 20 ± 3 °C with 50 ± 5% humidity and water and food available ad libitum. The animal study protocol was approved by the Institutional Animal Care and Use Committee at the University of Wisconsin-Madison.

### General methods for LYTACs synthesis and analysis

The small molecular chemical reaction was monitored by thin-layer chromatography and visualized with a UV lamp at 254 nm (MilliporeSigma). The polymers were synthesized by RAFT polymerization with Schlenk line (Sigma-Aldrich) for degassing. The small molecule inhibitors were linked to the polymers by amidation reaction. Nuclear magnetic resonance (NMR) spectra were collected using a Bruker Avance III HD 400-MHz NMR spectrometer and analyzed by MestReNova (version 14.0.0-23239, Mestrelab Research). The number-averaged molecular weight (Mn) of the polymer was determined by GPC using water or DMF as mobile phase. High-resolution mass spectrometry data were obtained using a Bruker MaXis Ultra-High Resolution Quadrupole Time-of-Flight mass spectrometer.

### Western blot

Western blot was used to investigate the expression of PD-L1, RAC1, Luciferase, or LC3B in MDA-MB-231 or 4T1-hPD-L1 cells after indicated treatments.

For the in vitro experiments, MDA-MB-231 cells were seeded in six-well culture plates at a density of 3 × 10^5^ cells in each well and cultured overnight. After treatments, cells were washed with PBS three times and lysed on ice with RIPA lysis buffer containing PMSF (1 mM) and a protease inhibitor cocktail (MedChemExpress, HY-K0011) for 30 min.

For the in vivo experiments, tumor tissues from 4T1-hPD-L1 tumor-bearing mice were collected and weighed, followed by being cut into small pieces. The tumor tissues were further incubated in collagenase A (Sigma-Aldrich, C5138)-contained DMEM solution (1 mg/mL) for 1.5 h at 37℃ and then ground to across 40 μm nylon mesh cell strainer (Falcon, 352340). The single-cell suspensions were centrifuged at 2,000 rpm for 8 min. The cell sediments were lysed on ice with RIPA lysis buffer for 30 min.

Lysates were collected and centrifuged at 14,000g for 10 min at 4 °C. The corresponding supernatants were collected, followed by quantification of total proteins in the supernatant using a bicinchoninic acid (BCA) assay kit (Thermo Fisher, 23225). Afterwards, the protein solution was mixed with loading buffer in a volume ratio of three to one and heated at 95 ℃ for 5 min. Samples (20 μg of total proteins per lane) were loaded into Bis-Tris gels formulated with Bis-Tris buffer, 30% acrylamide/Bis, 10% ammonium persulfate solution and TEMED. Bis-Tris 10% gels were used to analyze PD-L1, Luciferase and β-Actin and 15% gels were used to analyze LC-3B and RAC1. The electrophoresis voltage was 80 V for stacking gel and 130 V for resolving gel. Separated proteins were transferred to polyvinylidene fluoride membranes which were then blocked with 5% nonfat milk in PBST (0.5% Tween-20 in PBS). The membranes were incubated with anti-PD-L1 (1:1000, Cell Signaling Technology 13684), anti-Firefly Luciferase (1:1000, Cell Signaling Technology 25039), anti-RAC1 (1:1000, Cell Signaling Technology 4651) or anti-LC-3B (1:1000, Cell Signaling Technology 83506) primary antibody overnight at 4 °C, followed by incubation with goat anti-rabbit IgG H&L (HRP) secondary antibody at room temperature for 1 h. Membranes were treated with electrochemiluminescence western blot substrate (Thermo Fisher, 34578) and detected using the Odyssey XF Imaging System (LI-COR).

### Confocal imaging

Confocal imaging was carried out to visualize the localization of PD-L1, MIF, organelles and McR-TACs or autophagy level in MDA-MB-231 cells.

Live cells were used for MIF degradation, McR-TACs and autophagosomes imaging. MDA-MB-231 cells were seeded in 35 mm glass bottom dish (Cellvis, D35-20-1.5-N) at a density of 3 × 10^4^ cells in each well and cultured overnight. After treatments, cells were washed and incubated with different dyes for indicated times. Leica SP8 Confocal WLL STED Microscope was applied to observe the fluorescent target.

Fixed cells were used for PD-L1 degradation imaging. After treatments, cells were washed and fixed with cold 4% formaldehyde (Spectrum, F-260) for 15 min at room temperature. The fixed cells were then permeabilized with 0.5% Triton X-100 in PBS for 20 min at room temperature, followed by blocking with 1% BSA solution for 1 h at room temperature. Primary antibodies and fluorescent-labeled secondary antibodies were applied to stain the cells for 16 h at 4 °C and 2 h at room temperature. After staining with Hoechst 33342, the cells were subjected to the Leica SP8 Confocal WLL STED Microscope for imaging.

### Flow cytometry analysis

Flow cytometry was used to analyze the cellular uptake of LYTACs or MIF and the CD45^+^ CD3^+^ CD8^+^ T cell infiltration in tumors of TNBC mice models.

For the in vitro experiment: Cells were seeded in 24-well culture plates at a density of 3 × 10^4^ cells in each well and cultured overnight. Cells were pre-treated with siRNA or incubated with different endocytosis inhibitors or transferred into a chamber with low temperature for 1 h. Then, sulfo-Cy5-labeled McR-TACs were added for 1 h-incubation. Afterwards, cells were washed and collected for flow cytometry analysis (Thermo Fisher, Attune NxT) by recording 10,000 cells.

For the in vivo experiment: The single-cell suspensions were obtained with the same method as mentioned in the western blot section. The cell numbers were counted with hemocytometer (Bulldoh-Bio, DHC-N420) and adjusted to a density of 1 × 10^7^/mL. The single-cell suspension was further blocked with 5% BSA for 20 min at room temperature and incubated with indicated primary antibodies for another 30 min at room temperature. Finally, the stained cells were analyzed using flow cytometry (Thermo Fisher, Attune NxT) by recording 100,000 cells.

### Intratumoral kinetics of LYTACs in vivo

4T1-hPD-L1-Luc cells (1 × 10^7^/mL) were suspended in PBS containing 50% Matrigel (Corning, 354234). Six-week-old female BALB/c mice were anesthetized and injected with 100 μL of cell suspension into the fourth mammary gland fat pads. After the tumor volumes reached about 100 mm^3^, the mice were anesthetized and i.p. injected with D-luciferin potassium salt (150 mg/kg) in 100 μL of PBS. The bioluminescence of tumor-bearing mice was detected using a Lago imaging system (Spectral Instruments Imaging, Bruker) and separated into different groups with similar bioluminescence signals. Free Cy5.5 or sulfo-Cy5.5-labeled LYTACs with or without Exo1 (150 μM) were i.t. injected into tumors with a dosage of 0.2 mg/kg Cy5.5 in 20 μL of PBS. The fluorescent intensity of Cy5.5 (exposure time = 30 s, excitation power = 30%) was monitored at indicated times and further analyzed using Aura software (Version 4.5.0).

### Evaluation of LYTACs’ efficacy and biosafety in TNBC mice model

4T1-hPD-L1 cells (1 × 10^7^/mL) were suspended in PBS containing 50% Matrigel (Corning, 354234). Six-week-old female BALB/c mice were anesthetized and injected with 100 μL of cell suspension into the fourth mammary gland fat pads. After the tumor volumes reached about 100 mm^3^, PBS, BMS-1166 or LYTACs were i.t. or i.v. injected in 20 μL of PBS once every four days for a total of two times. Among them, N-TACs_PD-L1_ was i.t. injected after dispersing in a formulation of 10% DMSO, 40% polyethylene glycol 300, 5% Tween-80 and 45% PBS.

During the treatment timeline, body weight was monitored every two days. The tumor weight was measured on day eight. Tumors were collected for flow cytometry, western blot and cytokines analysis on day eight. Hematological and blood chemical analyses for treated postsurgical mice were performed after one week of the last treatment.

### Statistical analysis and reproducibility

Biological replicates were obtained from distinct samples rather than from repeated measurements of the same sample. For immunoblotting, the blots shown are representative of at least three biologically independent replicates with similar results. The confocal images and flow cytometry results shown are representative of three biologically independent replicates with similar results. Investigators were blinded to group allocation throughout both the experimental procedures and outcome assessment. Statistical analysis was performed using Graph Pad Prism software 8, Microsoft Excel (Version 2108), ImageJ (Version 1.51j8), Aura (Version 4.5.0) and FlowJo (Version 10.0). In the statistical analysis for comparison among multiple data groups, one-way analysis of variance (ANOVA) followed by a Tukey post hoc test was conducted. Two-way ANOVAs were performed for multiple comparisons in fluorescent signal intensity in vivo analysis. *P* values < 0.05 were considered statistically significant. Except for the data regarding fluorescent intensity of LYTACs in tumor, which are presented as mean ± s.e.m., all other values and error bars are reported as mean ± s.d.

## Reporting summary

Further information on research design is available in the Nature Portfolio Reporting Summary linked to this article.

## Data availability

All data supporting the findings of this study are available within the paper and Supplementary Information. Source data are provided with this paper.

## Acknowledgements

We want to thank the optical imaging core, small animal facilities, flow cytometry core, histological core, and UW-Madison Wisconsin Centers for Nanoscale Technology for their help with this study. This work was supported, in part, by METAVIVOR Foundation Early Career Research Grant Award, the American Cancer Society Research Scholar Grant (Grant number: RSG-23-1140821-01-ET, to Q.H.), and the V Foundation Scholar Grant (to Q.H.). We also thank the support from the University of Wisconsin Carbone Cancer Center Research Collaborative and the Pancreas Cancer Task Force, and the start-up package from the University of Wisconsin-Madison.

## Author contributions

P.L. and Q.H. conceived the project. P.L., Y.L., Y.Y., Y.C., and M.C. performed experiments and interpreted data. P.L. and Q.H. wrote the manuscript with input from all authors. Q.H. provided supervision.

---

**Extended Data Fig. 1:**
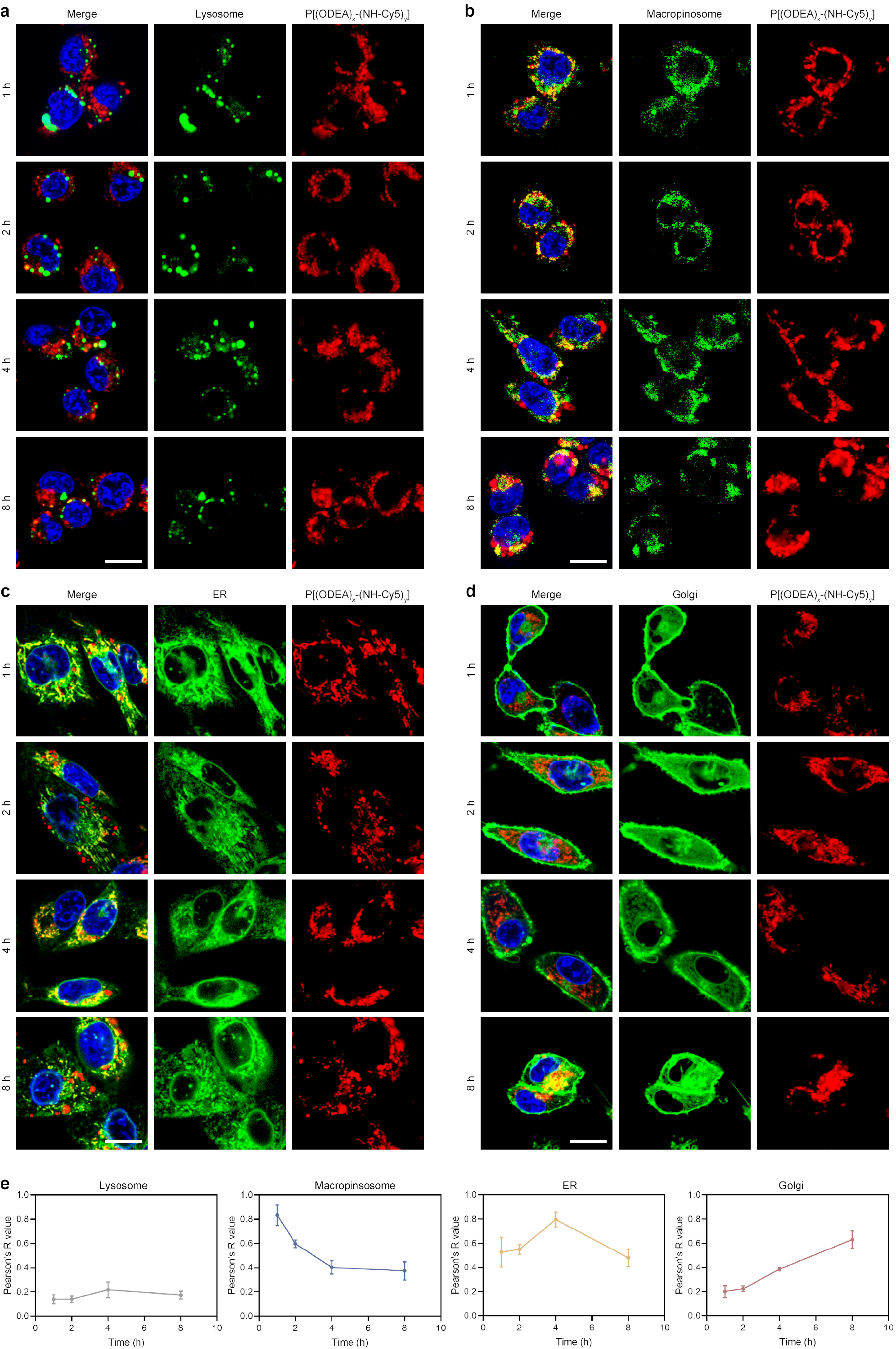
Intracellular transport process of zwitterionic polymers independent of warheads. **a-d**, Confocal imaging of the co-localization of P[(ODEA)_x_-(NH-Cy5)_y_] and organelles including lysosomes labeled with lysotracker green (**a**), macropinosomes labeled with TRITC-dextran (**b**), ER labeled with ER tracker green (**c**) and Golgi labeled with TMR ceramide (**d**) at indicated times. Scale bar, 30 μm. **e**, Pearson’s R value of confocal images from **a-d** (*n* = 3 per group). Data are presented as the mean values ± s.d.

**Extended Data Fig. 2:**
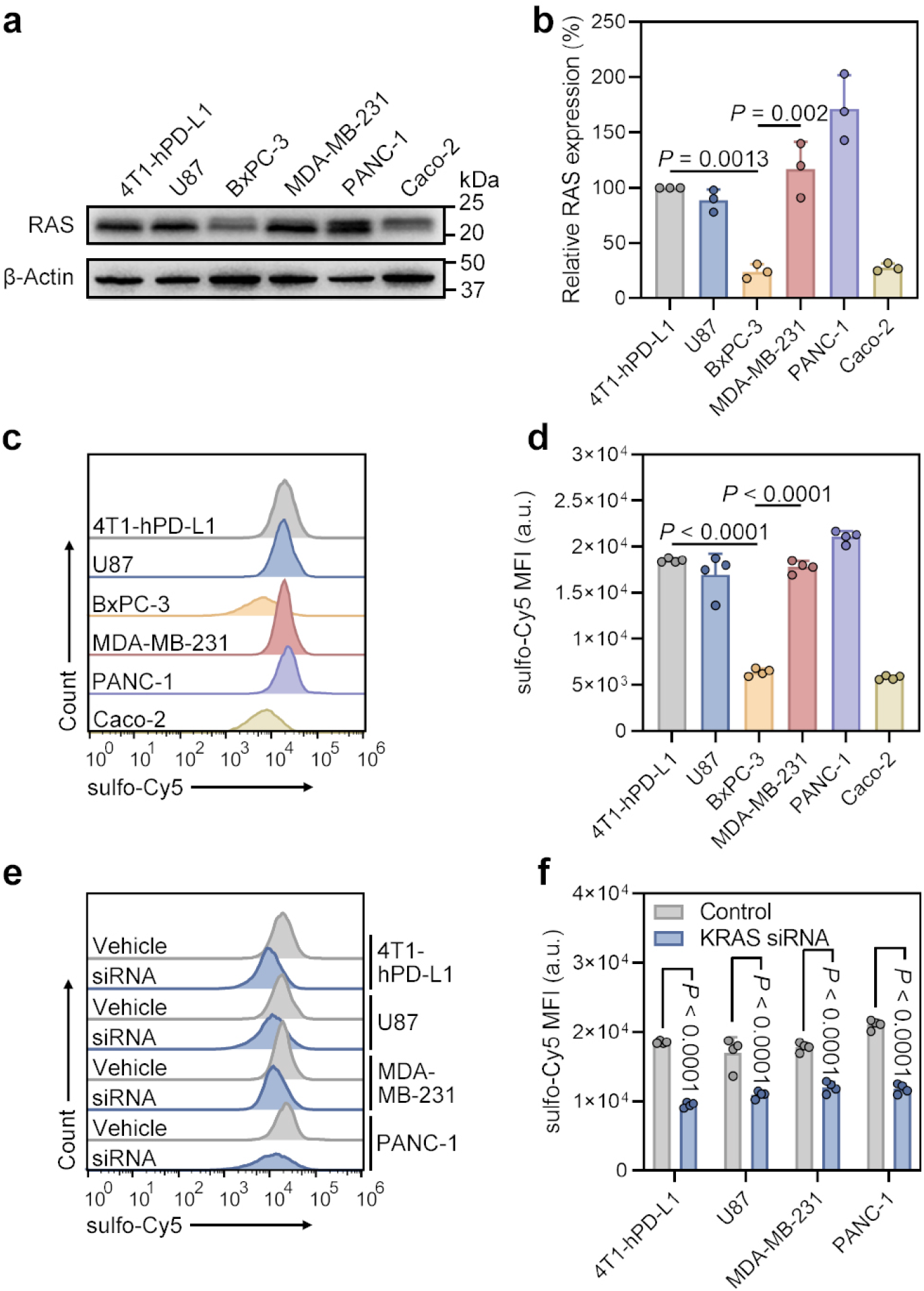
The internalization of McR-TACs_PD-L1_ in different cell lines positively correlates with RAS expression levels. **a-b**, Western blot (**a**) and semi-quantified analysis (**b**) of RAS levels in 4T1-hPD-L1, U87, BxPC-3, MDA-MB-231, PANC-1 and Caco-2 cells (*n* = 3 per group). **c-d**, Flow cytometry plot (**c**) and quantitative analysis (**d**) of sulfo-Cy5-McR-TACs_PD-L1_ internalization in above cell lines (*n* = 4 per group). **e-f**, Flow cytometry plot (**e**) and quantitative analysis (**f**) of sulfo-Cy5-McR-TACs_PD-L1_ internalization in 4T1-hPD-L1, U87, MDA-MB-231 and PANC-1 with or without KRAS knockdown (*n* = 4 per group). For **b** and **d**, data are presented as the mean values ± s.d. and were analyzed by a one-way ANOVA, followed by a Tukey’s multiple comparisons test. For **f**, data are presented as the mean values ± s.d. and were analyzed by two-tailed unpaired t-test.

**Extended Data Fig. 3:**
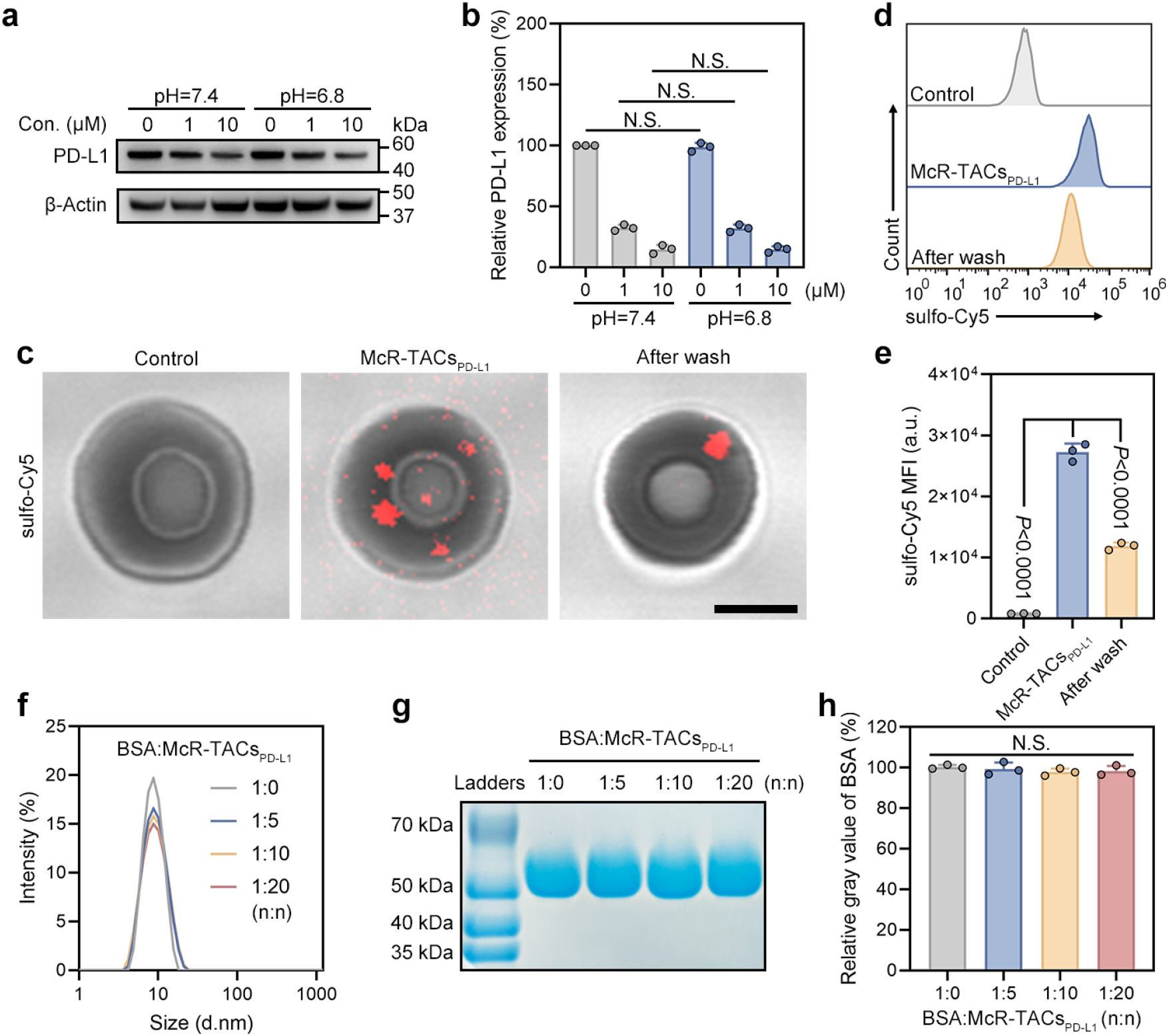
The influence of extracellular context on McR-TACs’ performance. **a-b**, Western blot (**a**) and semi-quantified analysis (**b**) of PD-L1 degradation in MDA-MB-231 cells by McR-TACs_PD-L1_ under different pH conditions (*n* = 3 per group). Data are presented as the mean values ± s.d. and were analyzed by a two-way ANOVA, followed by a Tukey’s multiple comparisons test. Not significant (N.S.). **c**, Confocal images of red blood cells incubated with sulfo-Cy5-McR-TACs_PD-L1_ before and after washout. Scale bar, 3 μm. **d-e**, Flow cytometry plot (**d**) and quantitative analysis (**e**) of sulfo-Cy5-McR-TACs_PD-L1_ adsorption on red blood cells before and after washout (*n* = 3 per group). **f**, The size distribution of BSA mixed with McR-TACs_PD-L1_ at different molar ratios. **g-h**, SDS-PAGE band (**g**) and semi-quantified analysis (**h**) of BSA molecular weight after mixed with McR-TACsPD-L1 at different molar ratios (*n* = 3 per group). For **e** and **h**, data are presented as the mean values ± s.d. and were analyzed by a one-way ANOVA, followed by a Tukey’s multiple comparisons test. Not significant (N.S.).

**Extended Data Fig. 4:**
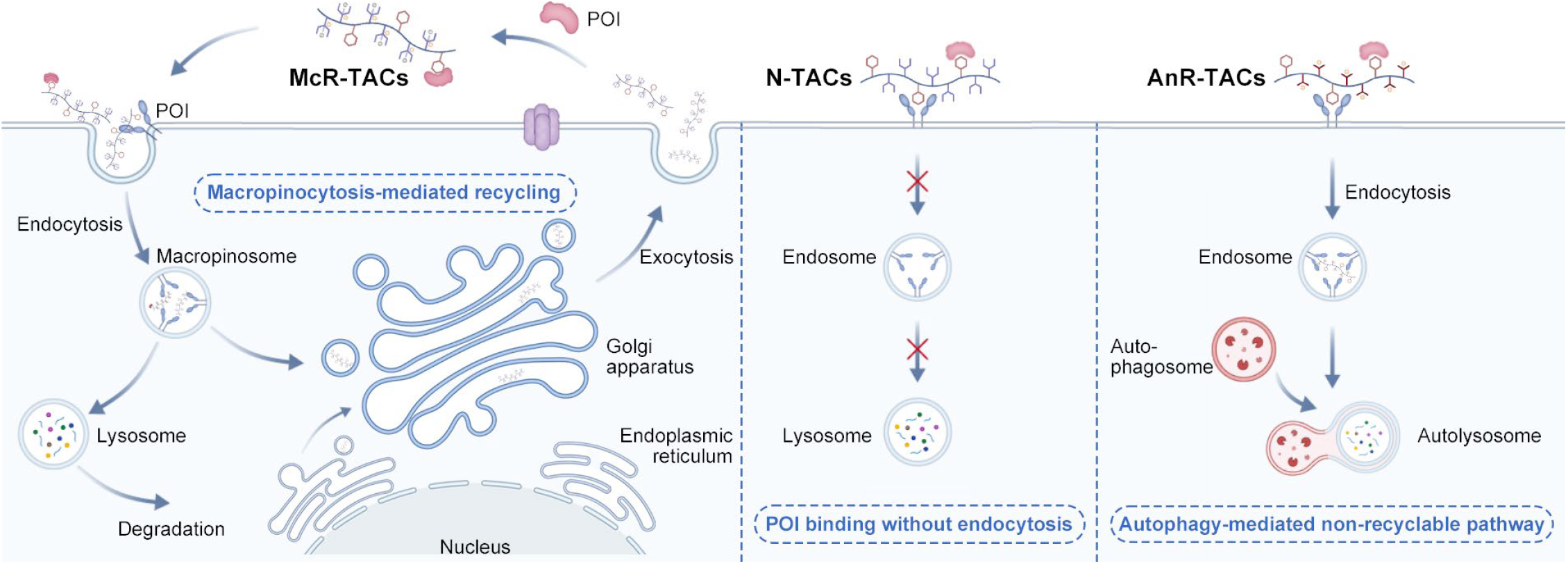
Schematic illustration of the intracellular processes and protein degradation mechanisms of McR-TACs, N-TACs and AnR-TACs. McR-TACs are recyclable LYTACs which could degrade both membrane and extracellular proteins through the macropinsosome-mediated transcytosis pathway. N-TACs could only bind to the POI without degrading them. AnR-TACs are non-recyclable LYTACs which could degrade both membrane and extracellular proteins through the autophagy pathway.

