## Supplemental files for "Macropinocytosis-Mediated Recyclable LYTACs (McR-TACs) for Membrane and Extracellular Protein Degradation"

### Table of Contents

1. Supplementary methods

2. Supplementary figures

3. Supplementary tables

4. Supplementary references

### 1. Supplementary methods

#### 1.1 Synthesis of McR-TACs$_{PD-L1}$ (x=~80, y=~5) (Fig. S1)[1]

(1) To get P[(DEA)$_x$-(NH-BOC)$_y$]: 4-cyano-4-(phenylcarbonothioylthio)pentanoic acid (CPPA, 20.0 mg, 0.072 mmol), 2-(N,N-diethylamino)ethyl methacrylate (DEA, 1061.0 mg, 5.73 mmol), 2-((tert-Butoxycarbonyl)amino)ethyl methacrylate (NH-BOC, 82.0 mg, 0.36 mmol) and 2,2'-azobis(2-methylpropionitrile) (AIBN, 11.8 mg, 0.072 mmol) were dissolved in anhydrous MeOH (4 mL) and added in a Schlenk tube. The solution was degassed through five repeated freeze-pump-thaw cycles with liquid nitrogen. The polymerization was carried out at 70 °C overnight. The reaction was quenched by opening the tube after completion, concentrated under reduced pressure, and precipitated with cold n-hexane for 3 times to give P[(DEA)$_x$-(NH-BOC)$_y$] in the form of salmon-colored viscous liquid (875.0 mg, yield 75.2%). $^1$H-NMR (400 MHz, Methanol-$d_4$) δ 4.10 (s, 169H), 2.83 (s, 161H), 2.68 (s, 320H), 2.12 – 1.83 (m, 350H), 1.52 (d, $J$ = 9.2 Hz, 142H), 1.19 – 1.07 (m, 664H), 1.03 – 0.74 (m, 462H).

(2) To get P[(ODEA)$_x$-(NH-BOC)$_y$]: P[(DEA)$_x$-(NH-BOC)$_y$] (800.0 mg) was dissolved in 5 mL 30% $H_2O_2$ solution and shaken at 100 rpm under room temperature for 4 h. The solution was then dialyzed against deionized water (3.5k MWCO) for 24 h and further lyophilized to get P[(ODEA)$_x$-(NH-BOC)$_y$] in the form of flocculent white solid (784.2 mg, yield 92.5%). $^1$H-NMR (400 MHz, Methanol- $d_4$) δ 4.50 (s, 160H), 3.58 (s, 166H), 3.39 (s, 312H), 2.14 – 1.79 (m, 357H), 1.51 (s, 106H), 1.35 (d, $J$ = 26.0 Hz, 593H), 1.20 – 0.77 (m, 622H).

(3) To get P[(ODEA)$_x$-(NH$_2$)$_y$]: P[(ODEA)$_x$-(NH-BOC)$_y$] (500.0 mg) was dissolved in 4 mL trifluoroacetic acid (TFA) and stirred at room temperature overnight. The reaction solution was further washed with dichloromethane (DCM) and concentrated in vacuo to remove TFA for five times. Then the solution was dialyzed against deionized water (3.5k MWCO) for 24 h and further lyophilized to get P[(ODEA)$_x$-(NH$_2$)$_y$] in the form of flocculent white solid (421.5 mg, yield 86.8%). $^1$H-NMR (400 MHz, Methanol-$d_4$) δ 4.51 (s, 160H), 3.78 (s, 175H), 3.62 – 3.46 (m, 305H), 2.14 – 1.79 (m, 346H), 1.42 (s, 516H), 1.21 – 0.81 (m, 555H).

(4) To get McR-TACs$_{PD-L1}$ (x=74, y=6): BMS-1166 (5.0 mg, 0.008 mmol), 1-hydroxybenzotriazole (HOBT, 2.1 mg, 0.016 mmol) and 1-(3-Dimethylaminopropyl)-3-ethylcarbodiimide (EDC, 2.4 mg, 0.016 mmol) were dissolved in anhydrous DMSO and stirred at room temperature for 15 min. Then P[(ODEA)$_x$-(NH$_2$)$_y$] (30.8 mg, 0.002 mmol) and triethylamine (TEA, 1.6 mg, 0.016 mmol) were added into the above reaction solution and stirred overnight. Then the solution was dialyzed against DMSO (3.5k MWCO) for 24 h and deionized water for 24 h and further lyophilized to get McR-TACs$_{PD-L1}$ in the form of flocculent white solid (24.9 mg, yield 72.4%). $^1$H-NMR (400 MHz, Methanol-$d_4$) δ 7.90 (s, 16H), 7.82 (d, $J$ = 8.0 Hz, 9H), 7.76 – 7.65 (m, 13H), 7.65 – 7.56 (m, 8H), 7.54 – 7.35 (m, 22H), 7.32 – 7.13 (m, 23H), 7.03 – 6.83 (m, 25H),

6.75 (dd, $J$ = 10.8, 1.8 Hz, 18H), 5.29 – 5.16 (m, 27H), 4.50 (s, 160H), 3.56 (d, $J$ = 12.9 Hz, 176H), 3.37 (s, 332H), 1.90 (s, 385H), 1.34 (d, $J$ = 27.7 Hz, 634H), 1.23 – 0.81 (m, 666H).

**1.2 Synthesis of McR-TACs$_{PD\text{-}L1}$ analogues**

The synthesis of McR-TACs$_{PD\text{-}L1}$ analogues (x=~10/20/40/80/120, y=~5) with different lengths is controlled by changing the dosage of DEA monomer. The synthetic procedures of other analogues are similar to those of McR-TACs$_{PD\text{-}L1}$ (x=~80, y=~5).

**1.3 Synthesis of McR-TACs$_{MIF}$ (x=~80, y=~5) (Fig. S9)**

(1) To get ISO-1-COOH: ISO-1 (50.0 mg, 0.213 mmol) was dissolved in cold MeOH and slowly added aqueous LiOH (25.0 mg, 1.063 mmol) over 15 min. The volume ratio of MeOH to water is 5 to 1. The reaction mixture was allowed to heat to 37 °C overnight with stirring. The organic solvent was removed in vacuo and the residual aqueous solution was acidified to pH 2 with 1N HCl. The aqueous phase was extracted with ethyl acetate for three times. The organic extract was dried over $MgSO_4$ and concentrated to get ISO-1-COOH in the form of white solid (34 mg, yield 72.3%). $^1$H-NMR (400 MHz, Methanol-$d_4$) δ 7.54 (d, $J$ = 8.7 Hz, 2H), 6.87 - 6.77 (m, 2H), 5.05 (dq, $J$ = 10.4, 7.1 Hz, 1H), 3.62 - 3.51 (m, 1H), 3.19 (ddd, $J$ = 16.9, 7.3, 2.2 Hz, 1H), 2.81 - 2.63 (m, 2H).

(2) To get McR-TACs$_{MIF}$ (x=68, y=10): ISO-1-COOH (5.0 mg, 0.023 mmol), HOBT (6.1 mg, 0.046 mmol) and EDC (7.0 mg, 0.046 mmol) were dissolved in anhydrous DMSO and stirred at room temperature for 15 min. Then P[(ODEA)$_x$-($NH_2$)$_y$] (87.4 mg, 0.005 mmol) and TEA (4.7 mg, 0.046 mmol) were added into the above reaction solution and stirred overnight. Then the solution was dialyzed against DMSO (3.5k MWCO) for 24 h and deionized water for 24 h and further lyophilized to get McR-TACs$_{MIF}$ in the form of flocculent white solid (67.5 mg, yield 73.3%). $^1$H-NMR (400 MHz, Methanol-$d_4$) δ 7.62 (s, 25H), 6.92 (s, 25H), 5.13 (s, 14H), 4.50 (s, 160H), 3.56 (s, 152H), 3.40 (s, 321H), 1.98 (d, $J$ = 57.0 Hz, 365H), 1.38 (s, 531H), 1.19 – 0.76 (m, 634H).

**1.4 Synthesis of N-TACs$_{PD\text{-}L1}$ (x=~80, y=~5) (Fig. S20)**

(1) To get P[(DEA)$_x$-($NH_2$)$_y$]: P[(DEA)$_x$-(NH-BOC)$_y$] (500.0 mg) was dissolved in 4 mL TFA and stirred at room temperature overnight. The reaction solution was further washed with DCM and concentrated in vacuo to remove TFA for five times. Then the solution was dialyzed against deionized water (3.5k MWCO) for 24 h and further lyophilized to get P[(DEA)$_x$-($NH_2$)$_y$] in the form of flocculent white solid (406.7 mg, yield 89.6%). $^1$H-NMR (400 MHz, Methanol-$d_4$) δ 4.11 (s, 160H), 2.86 (tt, $J$ = 12.3, 6.8 Hz, 181H), 2.80 – 2.62 (m, 317H), 1.94 (d, $J$ = 29.0 Hz, 315H), 1.11 (dd, $J$ = 23.6, 6.0 Hz, 656H), 1.03 – 0.70 (m, 445H).

(2) To get N-TACs$_{PD\text{-}L1}$ (x=80, y=4): BMS-1166 (5.0 mg, 0.008 mmol), HOBT (2.1 mg, 0.016 mmol) and EDC (2.4 mg, 0.016 mmol) were dissolved in anhydrous DMSO and stirred at room temperature for 15 min. Then P[(DEA)$_x$-($NH_2$)$_y$] (28.8

mg, 0.002 mmol) and TEA (1.6 mg, 0.016 mmol) were added into the above reaction solution and stirred overnight. Then the solution was dialyzed against DMSO (3.5k MWCO) for 24 h and deionized water for 24 h and further lyophilized to get N-TACs$_{PD-L1}$ in the form of flocculent white solid (24.9 mg, yield 72.4%). $^1$H-NMR (400 MHz, Methanol-$d_4$) 7.90 (s, 5H), 7.82 (d, $J$ = 8.0 Hz, 4H), 7.76 – 7.65 (m, 6H), 7.65 – 7.56 (m, 6H), 7.54 – 7.35 (m, 7H), 7.32 – 7.13 (m, 8H), 7.03 – 6.83 (m, 7H), 6.75 (dd, $J$ = 10.8, 1.8 Hz, 7H), 5.29 – 5.16 (m, 4H), 4.15 (s, 160H), δ 2.83 (s, 172H), 2.68 (s, 336H), 1.90 (s, 400H), 1.13 (s, 756H), 0.92 (s, 396H).

**1.5 Synthesis of AnR-TACs$_{PD-L1}$ (x=~80, y=~5) (Fig. S26)**

(1) To get P(NH-BOC)$_x$: CPPA (10.0 mg, 0.036 mmol), NH-BOC (656.5 mg, 2.880 mmol) and AIBN (5.9 mg, 0.036 mmol) were dissolved in anhydrous MeOH (4 mL) and added in a Schlenk tube. The solution was degassed through five repeated freeze-pump-thaw cycles with liquid nitrogen. The polymerization was carried out at 70 °C overnight. The reaction was quenched by opening the tube after completion, concentrated under reduced pressure, and precipitated with cold n-hexane for 3 times to give P(NH-BOC)$_x$ in the form of salmon thick liquid (589.3 mg, yield 88.6%). $^1$H-NMR (400 MHz, Methanol-$d_4$) δ 4.02 (s, 160H), 3.38 (s, 160H), 2.16 – 1.82 (m, 183H), 1.50 (s, 836H), 0.99 – 0.82 (m, 319H).

(2) To get P($NH_2$)$_x$: P(NH-BOC)$_x$ (500 mg) was dissolved in 4 mL TFA and stirred at room temperature overnight. The reaction solution was further washed with DCM and concentrated in vacuo to remove TFA for five times. Then the solution was dialyzed against deionized water (3.5k MWCO) for 24 h and further lyophilized to get P($NH_2$)$_x$ in the form of white solid (245.9 mg, yield 86.2%). $^1$H-NMR (400 MHz, Methanol-$d_4$) δ 4.02 (s, 160H), 3.37 (s, 164H), 1.96 (d, $J$ = 29.7 Hz, 169H), 0.95 (s, 189H).

(3) To get AnR-TACs$_{PD-L1}$ (x=71, y=5): BMS-1166 (5.0 mg, 0.008 mmol), HOBT (2.1 mg, 0.016 mmol) and EDC (2.4 mg, 0.016 mmol) were dissolved in anhydrous DMSO and stirred at room temperature for 15 min. Then P($NH_2$)$_x$ (17.0 mg, 0.002 mmol) and TEA (1.6 mg, 0.016 mmol) were added into the above reaction solution and stirred overnight. Then the solution was dialyzed against DMSO (3.5k MWCO) for 24 h and deionized water for 24 h and further lyophilized to get AnR-TACs$_{PD-L1}$ in the form of flocculent white solid (15.6 mg, yield 70.5%). $^1$H-NMR (400 MHz, Methanol-$d_4$) 7.90 (s, 14H), 7.82 (d, $J$ = 8.0 Hz, 20H), 7.76 – 7.65 (m, 8H), 7.65 – 7.56 (m, 14H), 7.54 – 7.35 (m, 12H), 7.32 – 7.13 (m, 15H), 7.03 – 6.83 (m, 14H), 6.75 (dd, $J$ = 10.8, 1.8 Hz, 15H), 5.29 – 5.16 (m, 27H), 4.16 (s, 160H), 3.15 (s, 169H), 1.99 (s, 169H), 0.96 (s, 160H).

**1.6 Synthesis of N-TACs$_{MIF}$ (x=~80, y=~5) and AnR-TACs$_{MIF}$ (x=~80, y=~5) (Fig. S35-38)**

(1) To get N-TACs$_{MIF}$ (x=65, y=10): ISO-1-COOH (5.0 mg, 0.023 mmol), HOBT (6.1 mg, 0.046 mmol) and EDC (7.0 mg, 0.046 mmol) were dissolved in anhydrous DMSO and stirred at room temperature for 15 min. Then P[(DEA)$_x$-($NH_2$)$_y$] (82.8

mg, 0.005 mmol) and TEA (4.7 mg, 0.046 mmol) were added into the above reaction solution and stirred overnight. Then the solution was dialyzed against DMSO (3.5k MWCO) for 24 h and deionized water for 24 h and further lyophilized to get N-TACs$_{MIF}$ in the form of flocculent white solid (61.1 mg, yield 79.2%). $^1$H-NMR (400 MHz, Methanol-$d_4$) δ 7.62 (s, 24H), 6.90 (s, 24H), 5.13 (s, 12H), 4.23 (s, 160H), 3.26 – 2.83 (m, 465H), 1.91 (s, 358H), 1.25 (s, 527H), 1.18 – 0.80 (m, 566H).

(2) To get AnR-TACs$_{MIF}$ (x=61, y=11): ISO-1-COOH (5.0 mg, 0.023 mmol), HOBT (6.1 mg, 0.046 mmol) and EDC (7.0 mg, 0.046 mmol) were dissolved in anhydrous DMSO and stirred at room temperature for 15 min. Then $P(NH_2)_x$ (48.9 mg, 0.005 mmol) and TEA (4.7 mg, 0.046 mmol) were added into the above reaction solution and stirred overnight. Then the solution was dialyzed against DMSO (3.5k MWCO) for 24 h and deionized water for 24 h and further lyophilized to get AnR-TACs$_{MIF}$ in the form of flocculent white solid (48.4 mg, yield 90.5%). $^1$H-NMR (400 MHz, Methanol-$d_4$) δ 7.57 (s, 36H), 6.89 (s, 36H), 5.10 (s, 18H), 4.20 (s, 156H), 3.23 (s, 158H), 1.99 (s, 165H), 1.05 (d, $J$ = 77.1 Hz, 278H).

**1.7 GPC analysis of LYTACs**

The hydrophilic polymers, such as McR-TACs and AnR-TACs, were dissolved in water (5 mg/mL) and injected into Ultrahydrogel 250 column (7.8 × 300 mm) for GPC analysis. The hydrophobic polymers, such as N-TAC, were dissolved in DMF (5 mg/mL) and injected into Jordi Gel DVB Mixed Bed column (250 mm). Standard PEG polymers with different molecular weights were used to fit the standard curve. The injection volume was 20 μL per sample and the system temperature was set as 40 °C. The Mn of samples was calculated by the standard curves according to the instructions.

**1.8 Fluorescent labeling of LYTACs**

(1) For the in vitro assays: McR-TACs (10.0 mg, 0.0005 mmol) and sulfo-Cyanine5 NHS ester (0.386 mg, 0.0005 mmol) were dissolved in 1 mL PBS 7.4 and stirred overnight at room temperature. The reaction solution was further dialyzed against PBS 7.4 for 24 h to remove free probe. $P[(ODEA)_x\text{-}(NH_2)_y]$ (8.4 mg, 0.0005 mmol) and Cyanine5 NHS ester (1.7 mg, 0.0025 mmol) were dissolved in 1 mL DMSO with the addition of 10 μL TEA and stirred overnight at room temperature. The reaction solution was further dialyzed against DMSO for 24 h and PBS 7.4 for 24 h to remove free probe.

(2) For the in vivo assays: McR-TACs$_{PD\text{-}L1}$ (10.0 mg, 0.0005 mmol) or AnR-TACs$_{PD\text{-}L1}$ (6.9 mg, 0.0005 mmol) and sulfo-Cyanine5.5 NHS ester (0.386 mg, 0.0005 mmol) were dissolved in 1 mL PBS 7.4 and stirred overnight at room temperature. The reaction solution was further dialyzed against PBS 7.4 for 24 h to remove free probe.

(3) Characterization of fluorescent labeled LYTACs: The excitation and emission spectrum of LYTACs were scanned by the microplate reader (CLARIOstar Plus, BMG LABTECH). The PD-L1 degradation effects of McR-TACs$_{PD\text{-}L1}$ with or without fluorescent label were analyzed by western blot.

**1.9 Fluorescent labeling of MIF and BSA**

MIF (100 μg, 0.008 μmol) or BSA (532 μg, 0.008 μmol) and sulfo-Cyanine5 NHS ester (6.2 μg, 0.008 μmol) or NHS-Fluorescein (38 μg, 0.008 μmol) were dissolved in 1 mL PBS 7.4 and stirred overnight at 4 °C. The reaction solution was further dialyzed against PBS 7.4 for 24 h at 4 °C to remove free probe.

**1.10 Intracellular transportation of zwitterionic polymers independent of warheads**

(1) MDA-MB-231 cells were treated with $P[(ODEA)_x\text{-}(NH\text{-}Cy5)_y]$ (Cy5 equivalent dose = 1.5 μg/mL) for 1 h and then changed into fresh medium for continued 1 h, 3 h and 7 h. Afterwards, cells were stained with lysotracker (1:500, Lumiprobe Corporation A1201), TRITC-Dextran (1 mg/mL, Thermo Fisher), ER tracker (1:1000, Lumiprobe Corporation 3891) or BDP® TMR ceramide (1:200, Lumiprobe Corporation 3085) with Hoechst 33342 (1:1000) for 30 min at 37 °C. Finally, cells were observed under Leica SP8 Confocal WLL STED Microscope.

(2) Colocalization analysis: The Coloc 2 plug-in of Image J was used to analyze the colocalization between organelles and polymers by Pearson's R value.

**1.11 Targeted protein degradation through lysosome in MDA-MB-231 cells**

(1) For PD-L1 imaging, 10 μM McR-$TACs_{PD\text{-}L1}$ or 2.5 μM BMS-1166 was used to treat the cells for 4 h. The fixed and blocked cells were incubated with anti-PD-L1 (1:100, Invitrogen PA5-20343) and anti-LAMP1 (1:100, Cell Signaling Technology 15665) antibodies overnight at 4 °C. Afterwards, cells were washed with PBS and incubated with anti-rabbit IgG (H+L), F(ab')2 Fragment (Alexa Fluor® 488 Conjugate) (1:1000, Cell Signaling Technology 4412) and anti-mouse IgG (H+L), F(ab')2 Fragment (Alexa Fluor® 594 Conjugate) (1:1000, Cell Signaling Technology 8890) for 2 h at room temperature, followed by staining with Hoechst 33342 (1:1000, Thermo Fisher 62249). Finally, the cells were subjected to the Leica SP8 Confocal WLL STED Microscope for imaging. Cells were also treated with BMS-1166 or BMS-8 at different concentrations and then subjected to western blot to verify the protein degradation effects of free warheads.

(2) For MIF imaging, 10 μM McR-$TACs_{MIF}$ or 50 μM ISO-1 with 0.5 μM sulfo-Cy5-MIF was used to treat the cells for 4 h. The live cells were incubated with lysotracker (1:500, Lumiprobe Corporation A1201) and wheat germ agglutinin (WGA)-rhodamine (1:500, Vector Laboratories RL-1022) for 30 min at 37 °C, followed by staining with Hoechst 33342 (1:1000). Finally, the cells were subjected to the Leica SP8 Confocal WLL STED Microscope for imaging.

**1.12 MIF internalization and degradation in vitro**

(1) MIF internalization: 10 μM McR-$TACs_{MIF}$ with 0.5 μM sulfo-Cy5-MIF was used to treat the cells for 1 h, 2 h, 4 h and 6 h. Afterwards, cells were collected and analyzed by flow cytometry. Every 10,000 cells was counted to determine sulfo-Cy5-positive cells at the RL1 channel.

(2) MIF degradation: 10 nM McR-$TACs_{MIF}$ and 1 nM sulfo-Cy5-MIF was added to the supernatant of 4T1-hPD-L1 cells.

Then, cells and supernatant were collected at indicated times for quantifying MIF concentration with ELISA. Cellular total protein was quantified with the BCA method and used as a reference. For the degradation pathway of MIF investigation, 10 nM McR-TACs$_{MIF}$ and 1 nM sulfo-Cy5-MIF were added to the supernatant of 4T1-hPD-L1 cells with or without CQ (100 μM) or Exo1 (50 μM) for 24 h. Afterwards, intracellular and extracellular MIF concentrations were quantified with ELISA.

**1.13 Cytotoxicity of warheads and LYTACs in vitro**

Different concentrations of warheads or LYTACs were added to the cell medium without serum for 24 h. Afterwards, cells were washed three times and incubated with 10% CCK-8 solution for 3 h. Finally, the absorbance of supernatant was determined by a microplate reader (CLARIOstar Plus, BMG LABTECH) and was further used to calculate the cell viability compared to the control group.

**1.13 Flow cytometry analysis of endocytosis inhibitor and RAC1 knockdown effects on cellular uptake**

Cells were pre-treated with different inhibitors, including filipin (7.5 μM), chlorpromazine (30 μM), EIPA (10 μM), dynasore (15 μM) for 1 h, and then sulfo-Cy5-labeled McR-TACs were added for 1 h incubation (sulfo-Cy5 equivalent dose = 0.5 μg/mL). For the construction of RAC1 knockdown cell line, RAC1 siRNA (100 nM) was mixed with Lipofectamine 2000 according to the instructions. MDA-MB-231 cells were incubated with siRNA-lipid complex in Opti-MEM medium for 6 h and then changed to normal culture medium for an additional 30 h. Negative siRNA and Lipofectamine 2000 reagent were used as a negative and vehicle control group, respectively. The RAC1 knockdown effect was verified by western blot. Energy-dependent uptake was performed in a 4 °C refrigerator. Afterwards, cells were washed twice with PBS and digested by trypsin, and then centrifuged at 1,200 rpm for 5 min to remove the trypsin. The collected cells were resuspended in 0.3 ml PBS and analyzed by flow cytometry. Every 10,000 cells was counted to determine sulfo-Cy5-positive cells at the RL1 channel.

**1.14 McR-TACs internalization in different cell lines**

4T1-hPD-L1 with or without KRAS knock down, U87 with or without KRAS knock down, BxPC-3, MDA-MB-231 with or without KRAS knock down, PANC-1 with or without KRAS knock down and Caco-2 cells were treated with sulfo-Cy5-labeled McR-TACs for 1 h (sulfo-Cy5 equivalent dose = 0.5 μg/mL). Then, cells were collected and analyzed by flow cytometry. Every 10,000 cells was counted to determine sulfo-Cy5-positive cells at the RL1 channel. The RAS expression levels of the above cell lines were determined by western blot.

**1.15 McR-TACs and POI distribution in MDA-MB-231 cells**

(1) For PD-L1 and McR-TACs$_{PD-L1}$ imaging: Cells were incubated with sulfo-Cy5-labeled McR-TACs$_{PD-L1}$ for 1 h (sulfo-Cy5 equivalent dose = 1.5 μg/mL) and then changed into fresh medium for 1 h, 3 h and 7 h. Afterwards, the cells were fixed,

blocked and then stained with anti-PD-L1 (1:100, Invitrogen PA5-20343) and anti-LAMP1 (1:100, Cell Signaling Technology 15665), anti-Calnexin (1:200, proteintech 66903) or anti-GOLGA2/GM130 (1:500, proteintech 66662) antibodies overnight at 4 °C. The macropinosome was labeled by Fluorescein-BSA added with sulfo-Cy5-McR-$TACs_{PD-L1}$ together. Afterwards, cells were washed with PBS and incubated with anti-rabbit IgG (H+L), F(ab')2 Fragment (Alexa Fluor® 488 Conjugate) (1:1000, Cell Signaling Technology 4412) and anti-mouse IgG (H+L), F(ab')2 Fragment (Alexa Fluor® 594 Conjugate) (1:1000, Cell Signaling Technology 8890) for 2 h at room temperature, followed by staining with Hoechst 33342 (1:1000, Thermo Fisher 62249). Finally, the cells were subjected to the Leica SP8 Confocal WLL STED Microscope for imaging.

(2) For MIF and McR-$TACs_{MIF}$ imaging: Cells were incubated with sulfo-Cy5-labeled McR-$TACs_{MIF}$ for 1 h (sulfo-Cy5 equivalent dose = 1.5 μg/mL) and then changed into fresh medium for 1 h, 3 h and 7 h. Afterwards, the cells were stained with different organelle fluorescent probes as mentioned above in section 1.10. Finally, the cells were subjected to the Leica SP8 Confocal WLL STED Microscope for imaging.

**1.16 Targeted protein degradation pathway investigation of McR-TACs in MDA-MB-231 cells**

(1) McR-$TACs_{PD-L1}$: Cells were incubated with 10 μM McR-$TACs_{PD-L1}$ with or without BMS-1166 (10 μM), EIPA (10 μM), CQ (100 μM) or Exo1 (50 μM) for 24 h at 37°C. Afterwards, cells were washed twice with PBS and digested by trypsin. The collected cells were lysed for western blot analysis.

(2) McR-$TACs_{MIF}$: sulfo-Cy5-labeled MIF (0.5 μM) and 10 μM McR-$TACs_{MIF}$ were added to MDA-MB-231 cells pretreated with ISO-1 (500 μM). EIPA (10 μM), negative siRNA (100 nM) or RAC1 siRNA (100 nM) for 2 h. Afterwards, cells were collected and subjected to flow cytometry. Every 10,000 cells was counted to determine sulfo-Cy5-positive cells at the RL1 channel.

**1.17 The effects of Exo1 treatment on the internalization of McR-TACs in MDA-MB-231 cells**

Cells were treated with sulfo-Cy5-labeled McR-$TACs_{PD-L1}$ or sulfo-Cy5-labeled McR-$TACs_{MIF}$ and Exo1 (50 μM) for 1 h incubation (sulfo-Cy5 equivalent dose = 0.5 μg/mL). Then cells were collected and analyzed by flow cytometry. Every 10,000 cells was counted to determine sulfo-Cy5-positive cells at the RL1 channel.

**1.18 Recyclable degradation of POI by McR-TACs in MDA-MB-231 cells[2]**

(1) PD-L1: Cells were incubated with 20 μM sulfo-Cy5-labeled McR-$TACs_{PD-L1}$ for 4 h. Afterwards, cells were washed and added fresh culture medium with or without Exo1 (50 μM) for further incubation. Then, cells were collected at different times for PD-L1 level analysis by western blot.

(2) MIF: Cells were incubated with 10 μM McR-$TACs_{MIF}$ and 0.5 μM sulfo-Cy5-MIF for 4 h. Afterwards, cells were washed

and added fresh culture medium with 0.5 μM Fluorescein-MIF for further incubation. Then, cells were collected at different times for sulfo-Cy5-MIF degradation and Fluorescein-MIF internalization analysis by flow cytometry.

**1.19 Autophagy activation by AnR-TACs$_{PD-L1}$ in MDA-MB-231 cells**

Cells were incubated with indicated concentrations of AnR-TACs$_{PD-L1}$ with or without Baf-A1 (0.1 μM) for 24 h. Afterwards, cells were washed twice with PBS and digested by trypsin, and then centrifuged at 1,200 rpm for 5 min to remove the trypsin. The collected cells were lysed for western blot analysis of PD-L1 degradation and LC-3B activation.

To observe the autophagosome number changes, cells were treated with different groups of LYTACs (10 μM) for 24 h and stained with acridine orange (1 μM) for 20 min at room temperature. Finally, the cells were subjected to the Leica SP8 Confocal WLL STED Microscope for imaging. Rapamycin (10 μM) was used as a positive control.

**1.20 Extracellular context interaction with McR-TACs$_{PD-L1}$**

(1) pH condition: Different concentrations of McR-TACs$_{PD-L1}$ were added to the supernatant with normal (7.4) and acid (6.8) pH for 24 h. Then, cells were collected and subjected to western blot for PD-L1 degradation analysis.

(2) Red blood cells adsorption: The whole blood was obtained from Balb/c mice and centrifuged at 1200 rpm to collect red blood cells. Red blood cell solution was washed with PBS three times and diluted into $1.0 \times 10^5$/mL. Sulfo-Cy5-McR-TACs$_{PD-L1}$ was added into the above solution (sulfo-Cy5 equivalent dose = 0.5 μg/mL) and incubated for 1 h at 37°C. Then cells were washed twice to remove free polymers and passed through a 22G (0.7 mm diameter) syringe 30 times, followed by washing once. Finally, the red blood cell suspension before and after washout was subjected to flow cytometry and confocal imaging to analyze the reversible adsorption of McR-TACs$_{PD-L1}$ to red blood cells.

(3) BSA adsorption: BSA solution (1 mg/mL) was mixed with McR-TACs$_{PD-L1}$ at molar ratios of 1:0, 1:5, 1:10, 1:20 and stirred for 2 h at 37°C. Free polymers were removed by dialysis for 24 h. DLS and SDS-PAGE were used to analyze the size distribution and molecular weight of BSA, respectively.

**1.21 $CD45^+$ $CD3^+$ $CD8^+$ T cells infiltration in tumors of TNBC mice model**

After single-cell suspensions were obtained, DAPI (1:1000) and specific primary antibodies were added to label living effector T cell markers, including FITC anti-mouse CD45 (1:100), APC anti-mouse CD3 (1:80) and PE anti-mouse CD8a (1:80).

**1.22 Tumor cytokines analysis in vivo**

The supernatant was collected from the single-cell suspensions after centrifugation. The concentrations of IFN-γ and TNF-α were determined according to the instructions of ELISA kits and normalized with the tumor weights.

**1.23 Extracellular protein degradation in vivo**

4T1-hPD-L1 cells ($1 \times 10^7$/mL) were suspended in PBS containing 50% Matrigel (Corning, 354234). Six-week-old female BALB/c mice were anesthetized and injected with 100 µL of cell suspension into the fourth mammary gland fat pads. After the tumor volumes reached about 100 $mm^3$, MIF (0.5 mg/kg) with or without ISO-1, LYTACs were i.t. or i.v. injected in 20 µL of PBS. After 12 h, tumors were collected and made into single-cell suspensions which were further centrifuged and lysed for MIF ELISA analysis. The total protein concentration of the cell lysate was determined by BCA assay kit for normalization.

**1.24 Biosafety analysis of LYTACs for PD-L1 degradation in vivo**

The whole blood was collected from the model mice and directly analyzed using an AbaxisHM5 Complete Blood Count Analyzer. Part of the whole blood was centrifuged (1,000 rpm) to get plasma at 4°C and analyzed using an alanine transaminase (ALT) colorimetric activity assay kit (Cayman Chemical, 700260), aspartate aminotransferase (AST) colorimetric activity assay kit (Cayman Chemical, 701640) and a urea (BUN) colorimetric assay kit (Elabscience, E-BC-K183-M) according to the instructions.

### 2. Supplementary figures

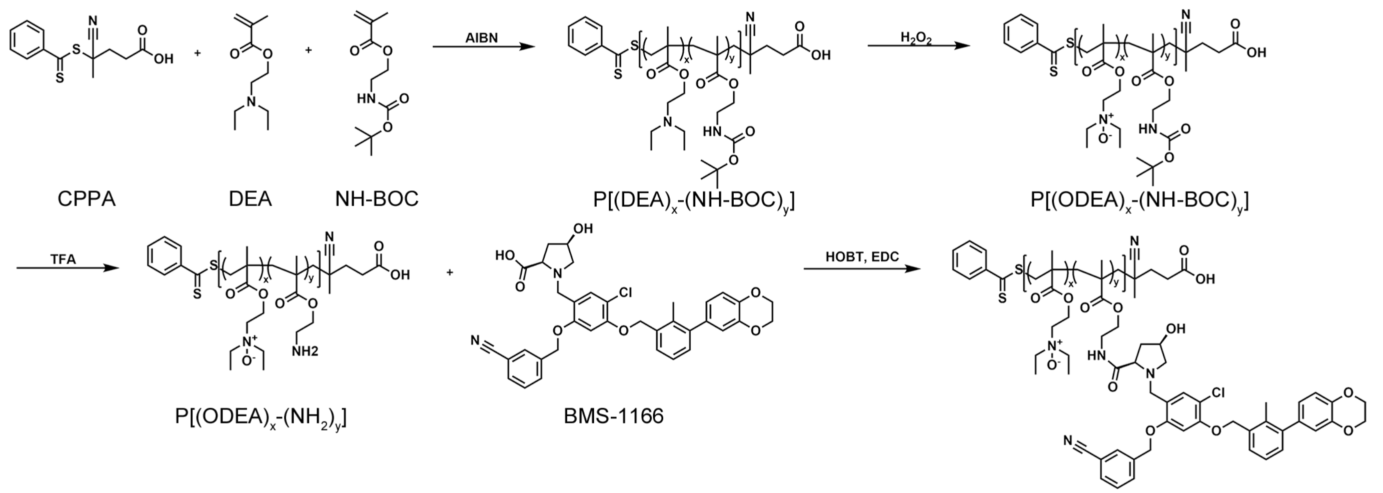


**Fig. S1**. Synthetic route of McR-TACs$_{PD-L1}$.

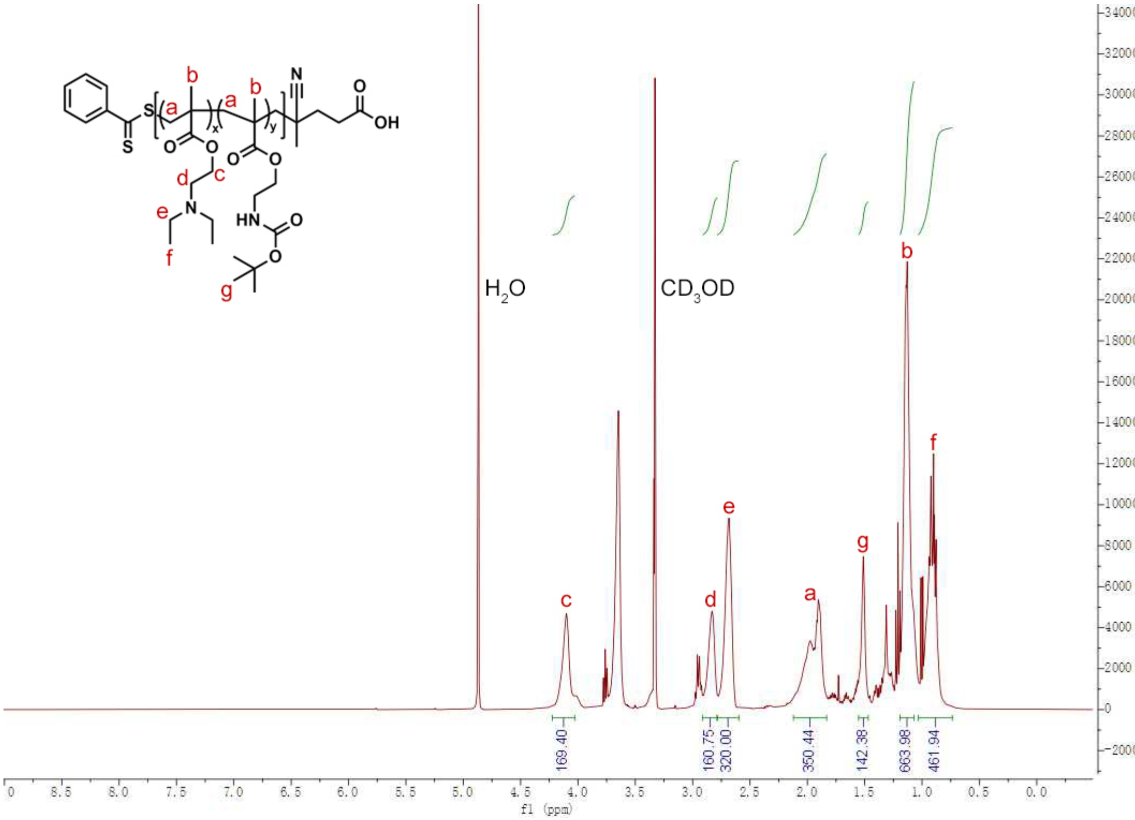


**Fig. S2**. $^{1}$H-NMR spectrum of P[(DEA)$_x$-(NH-BOC)$_y$] in $CD_3OD$.

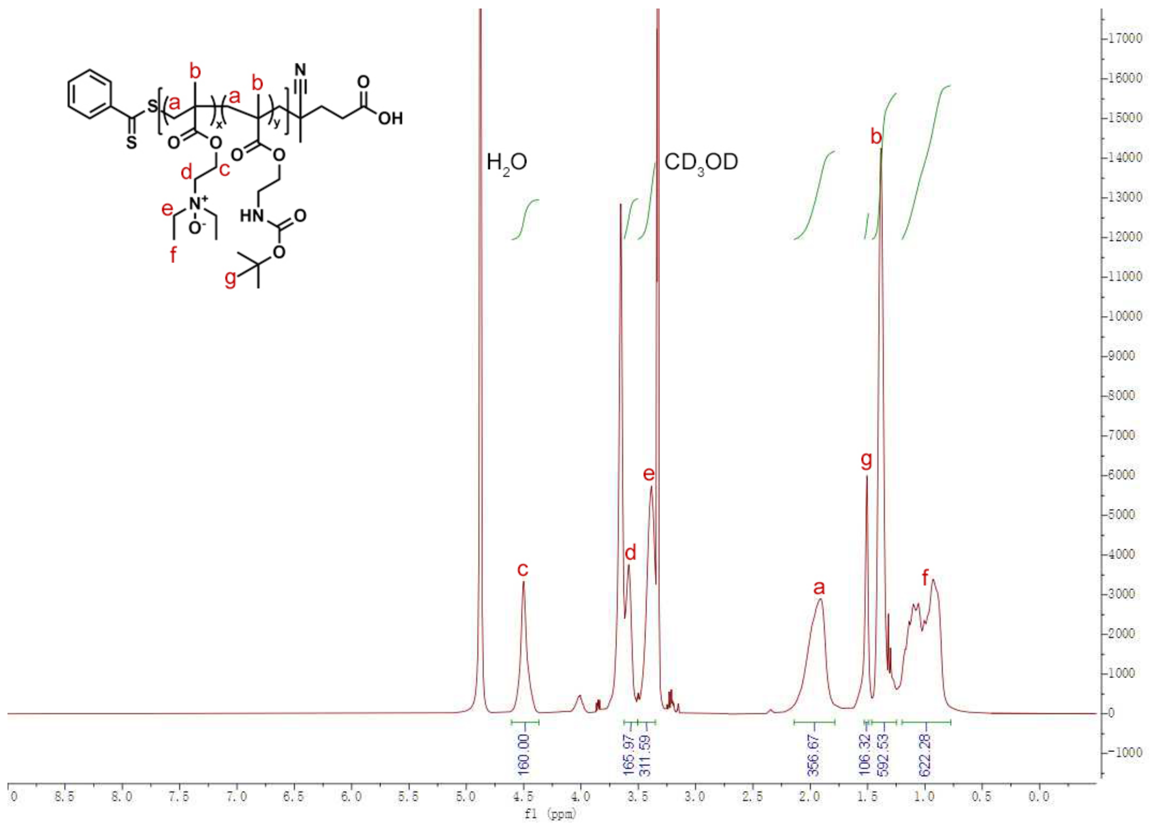


**Fig. S3**. $^{1}H$-NMR spectrum of P[$(ODEA)_x$-$(NH\text{-}BOC)_y$] in $CD_3OD$.

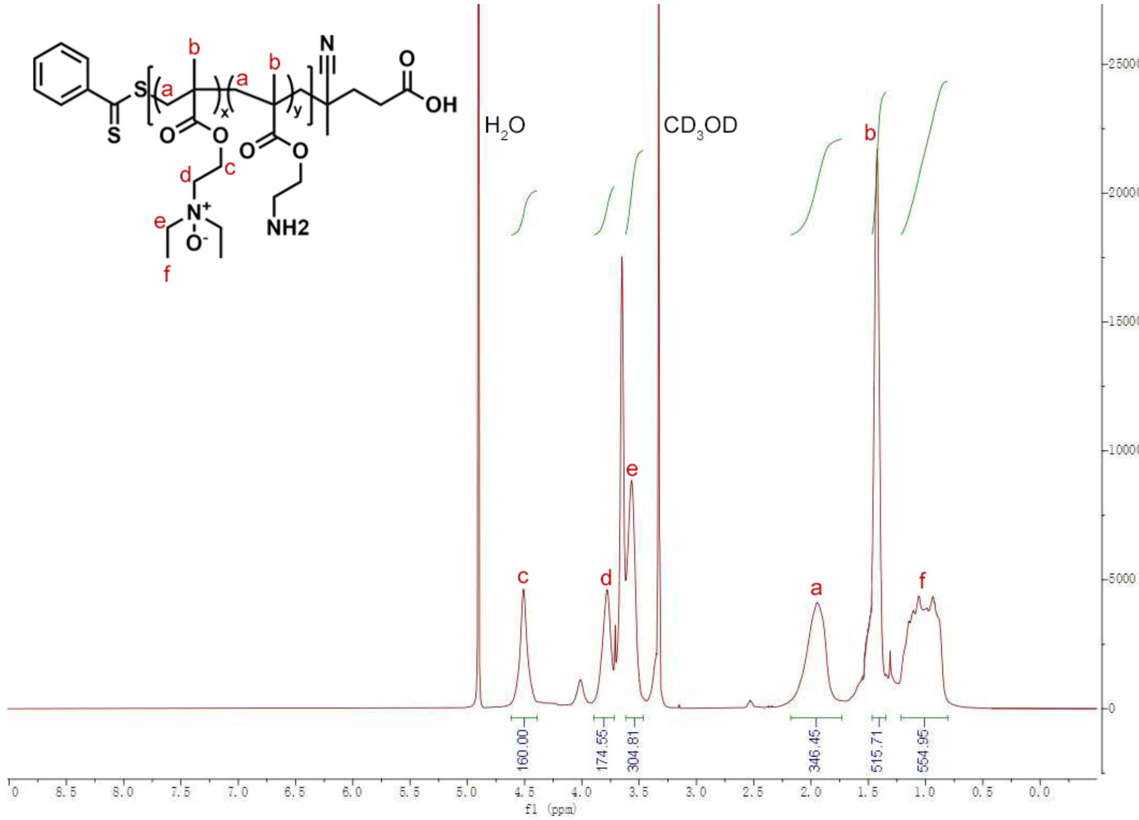


**Fig. S4**. $^1$H-NMR spectrum of $P[(ODEA)_x\text{-}(NH_2)_y]$ in $CD_3OD$.

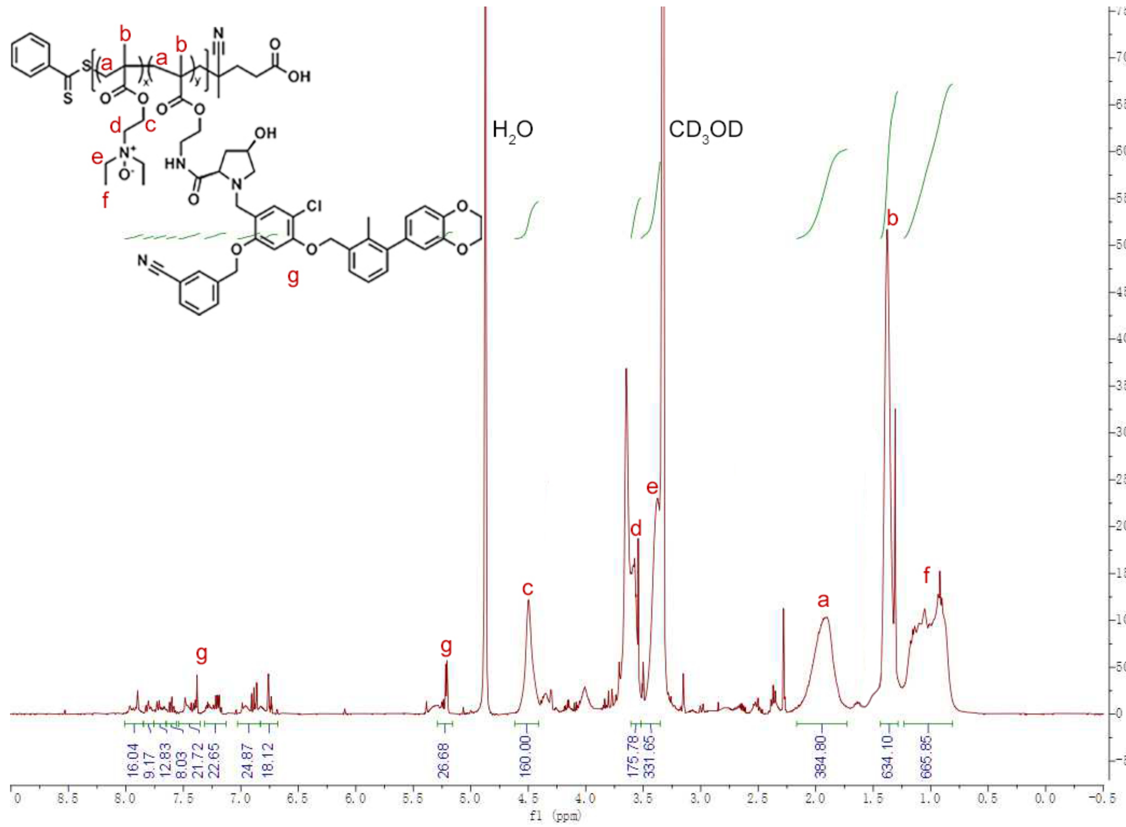


**Fig. S5**. $^1$H-NMR spectrum of McR-TACs$_{PD-L1}$ in $CD_3OD$.

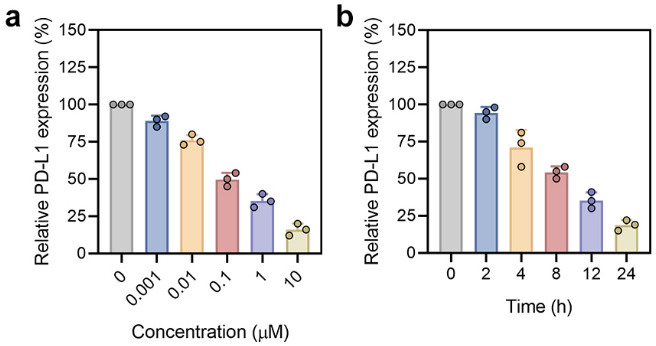


**Fig. S6**. Semi-quantified analysis of dose- (**a**) and time- (**b**) dependent PD-L1 degradation of McR-TACs$_{PD-L1}$ in MDA-MB-231 cells ($n$ = 3 per group). Data are presented as the mean values ± s.d.

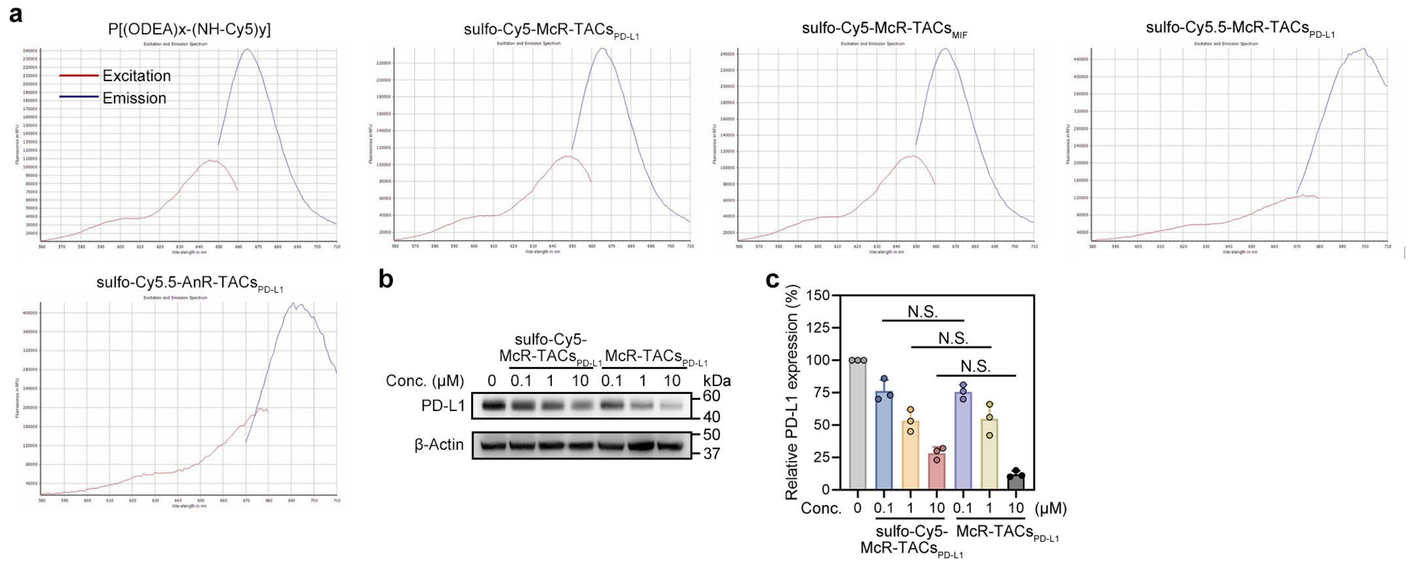


**Fig. S7**. **Characterization of fluorescent probe-labeled LYTACs. a**, Excitation and emission spectrum of fluorescent probe-labeled LYTACs. **b**, Western blot analysis of PD-L1 degradation in MDA-MB-231 cells treated with McR-TACs$_{PD-L1}$ with or without sulfo-Cy5 labeling under indicated concentrations for 24 h. **c**, Semi-quantification of relative PD-L1 levels in **b** ($n$ = 3 per group). Data are presented as the mean values ± s.d. and were analyzed by a two-way ANOVA, followed by a Tukey's multiple comparisons test. Not significant (N.S.).

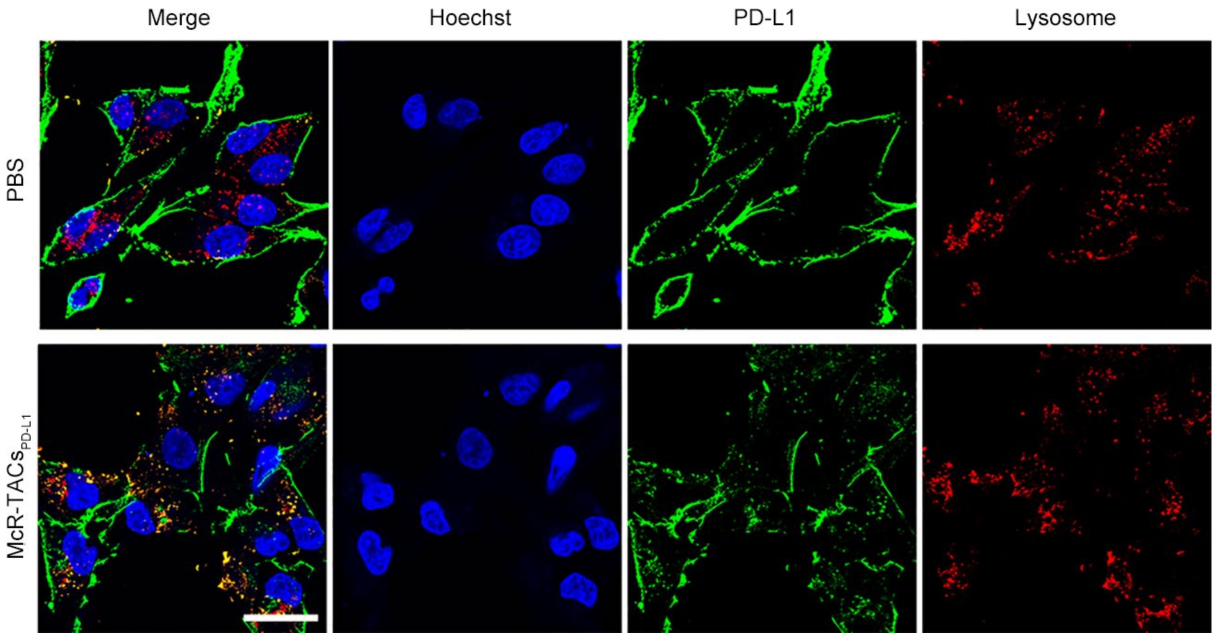


**Fig. S8**. Confocal imaging of the co-localization between PD-L1 and the lysosomes labeled anti-LAMP1 in MDA-MB-231 cells upon 8 h treatment with PBS or 10 μM McR-TACs$_{PD-L1}$. Scale bar, 40 μm.

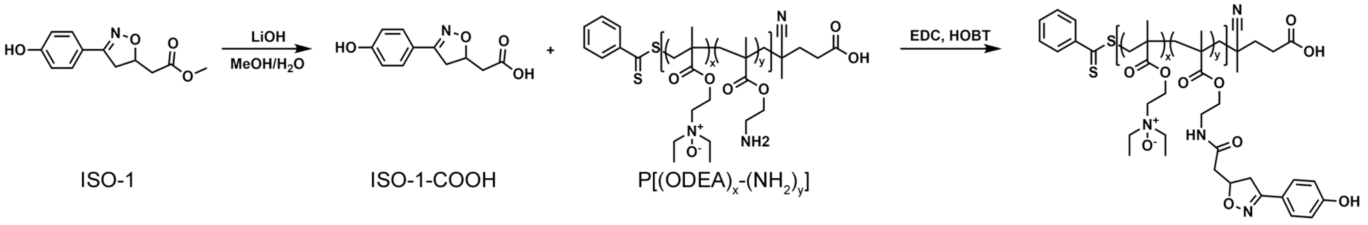


**Fig. S9**. Synthetic routes of McR-TACs$_{MIF}$.

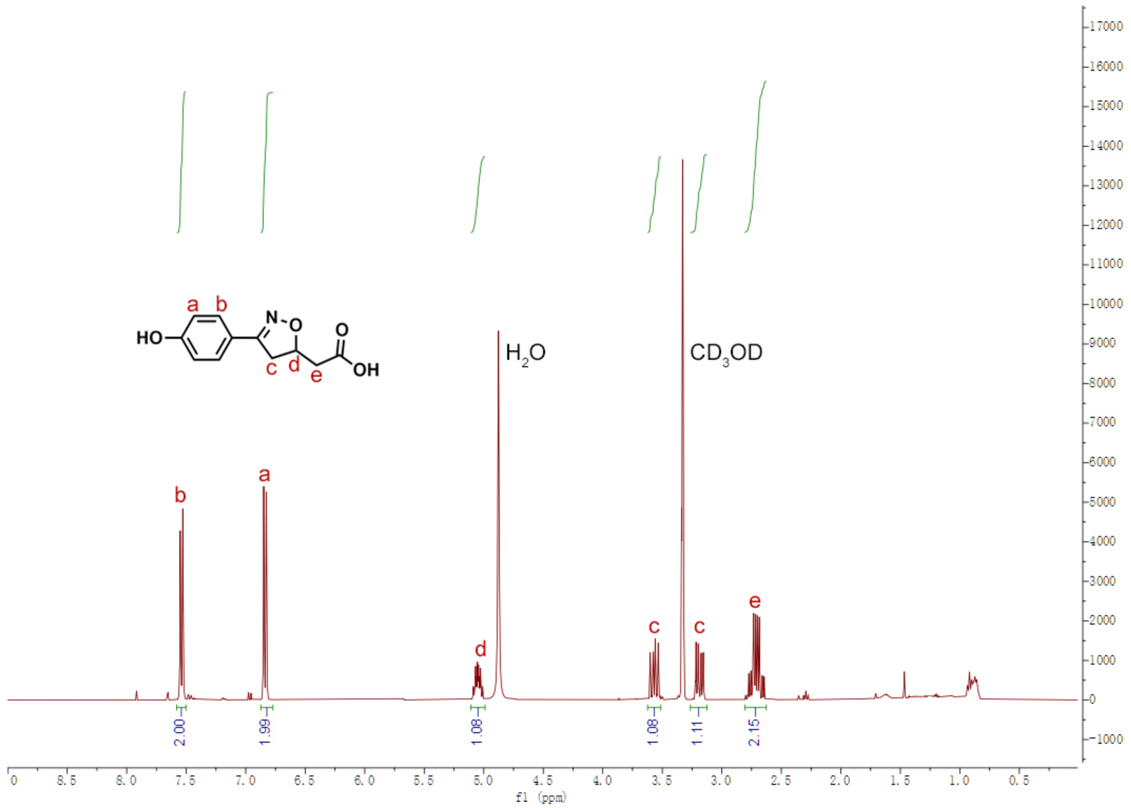


**Fig. S10**. ¹H-NMR spectrum of ISO-1-COOH in $CD_3OD$.

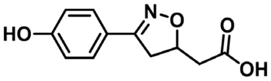


**Chemical Formula:** $C_{11}H_{11}NO_4$
**Exact Mass: 221.07**
**Molecular Weight: 221.21**
**m/z: 221.07 (100.0%), 222.07 (12.4%)**
**Elemental Analysis: C, 59.73; H, 5.01; N, 6.33; O, 28.93**

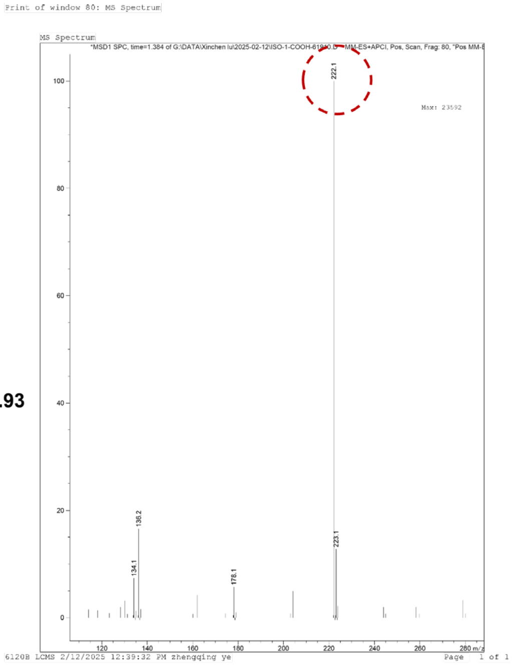


**Fig. S11**. MS spectrum of ISO-1-COOH.

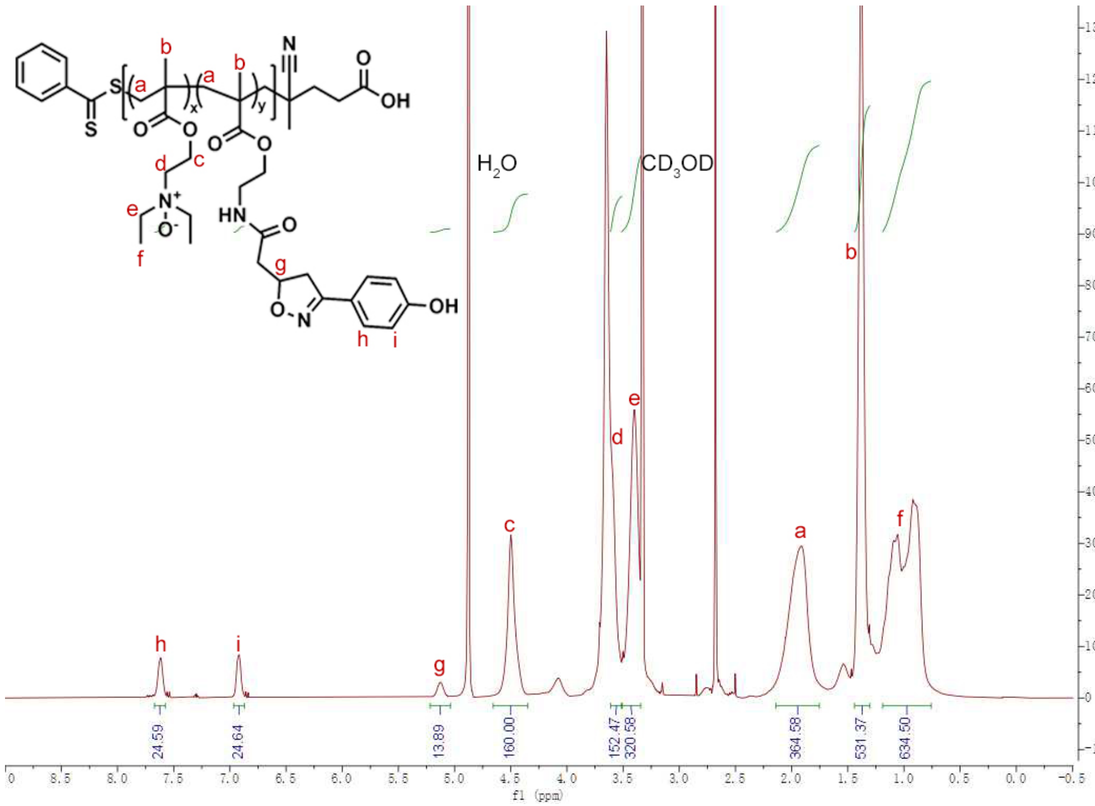


**Fig. S12**. [1]H-NMR spectrum of McR-TACs$_{MIF}$ in $CD_3OD$.

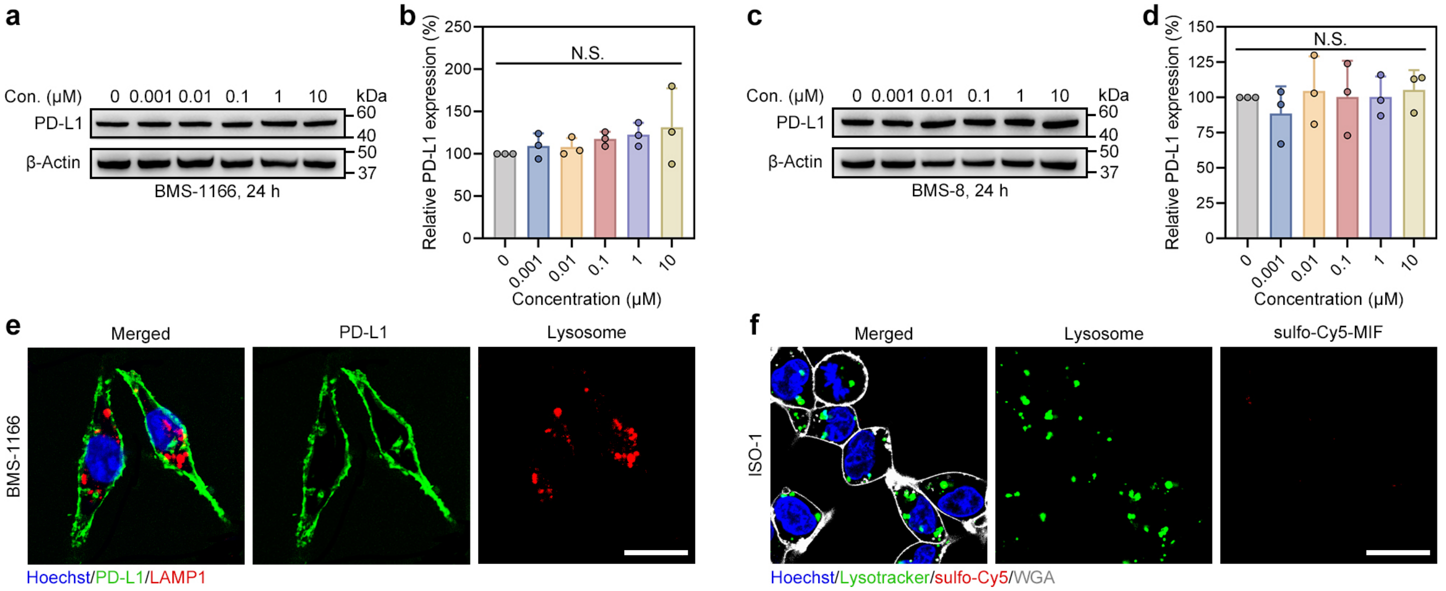


**Fig. S13**. **The influence of warheads on protein degradation. a-d**, Western blot analysis of PD-L1 degradation in MDA-MB-231 cells treated with BMS-1166 (**a**) or BMS-8 (**c**) under indicated concentrations for 24 h. Semi-quantification (**b**, **d**) of relative PD-L1 level in **a** and **c** ($n$ = 3 per group), respectively. Data are presented as the mean values ± s.d. and were analyzed by a one-way ANOVA, followed by a Tukey's multiple comparisons test. Not significant (N.S.). **e-f**, Confocal imaging of the localization of PD-L1 (**e**) or MIF (**f**) and lysosomes after warhead alone treatment. Scale bar, 40 μm.

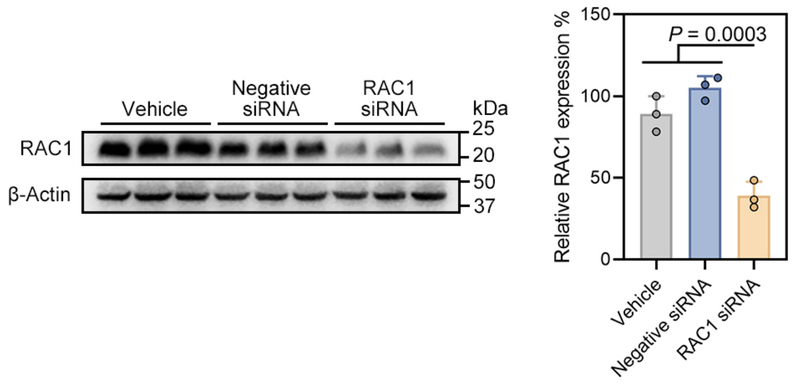


**Fig. S14**. Western blot analysis of RAC1 levels in MDA-MB-231 cells treated with vehicle, negative siRNA and RAC1 siRNA. Semi-quantification of relative RAC1 level in blots ($n$ = 3 per group). Data are presented as the mean values ± s.d.

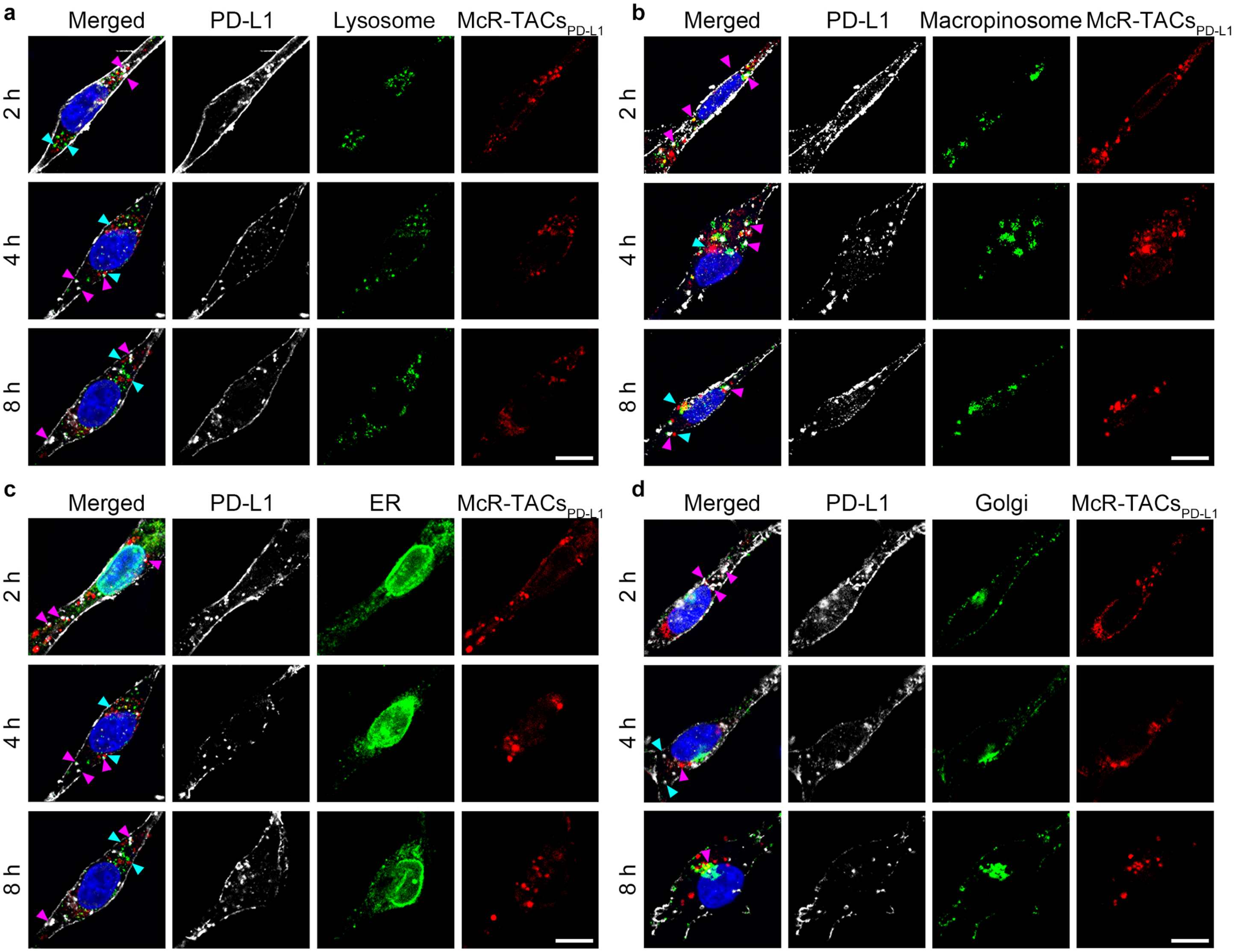


**Fig. S15**. Confocal imaging of the localization of sulfo-Cy5-McR-TACs$_{PD-L1}$, PD-L1 and organelles including lysosomes stained with anti-LAMP1 (**a**), macropinosomes labeled with Fluorescein-BSA (**b**), ER stained with anti-Calnexin (**c**) and Golgi stained with anti-GM130 (**d**) at indicated times. The purple arrowheads point to the co-localized signals and the blue arrowheads point to the independent signals. Scale bar, 20 μm.

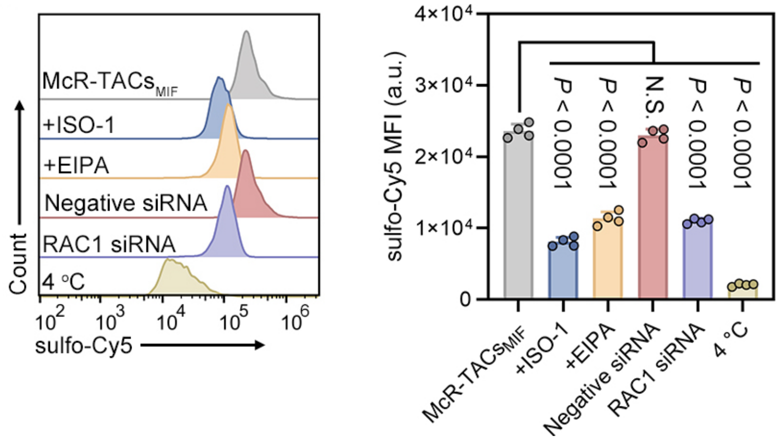


**Fig. S16**. Flow cytometry analysis of sulfo-Cy5-MIF internalization mediated by McR-TACs$_{MIF}$ in MDA-MB-231 cells pre-treated with indicated inhibitory conditions ($n = 4$ per group). Data are presented as the mean values ± s.d. and were analyzed by a one-way ANOVA, followed by a Tukey's multiple comparisons test. Not significant (N.S.).

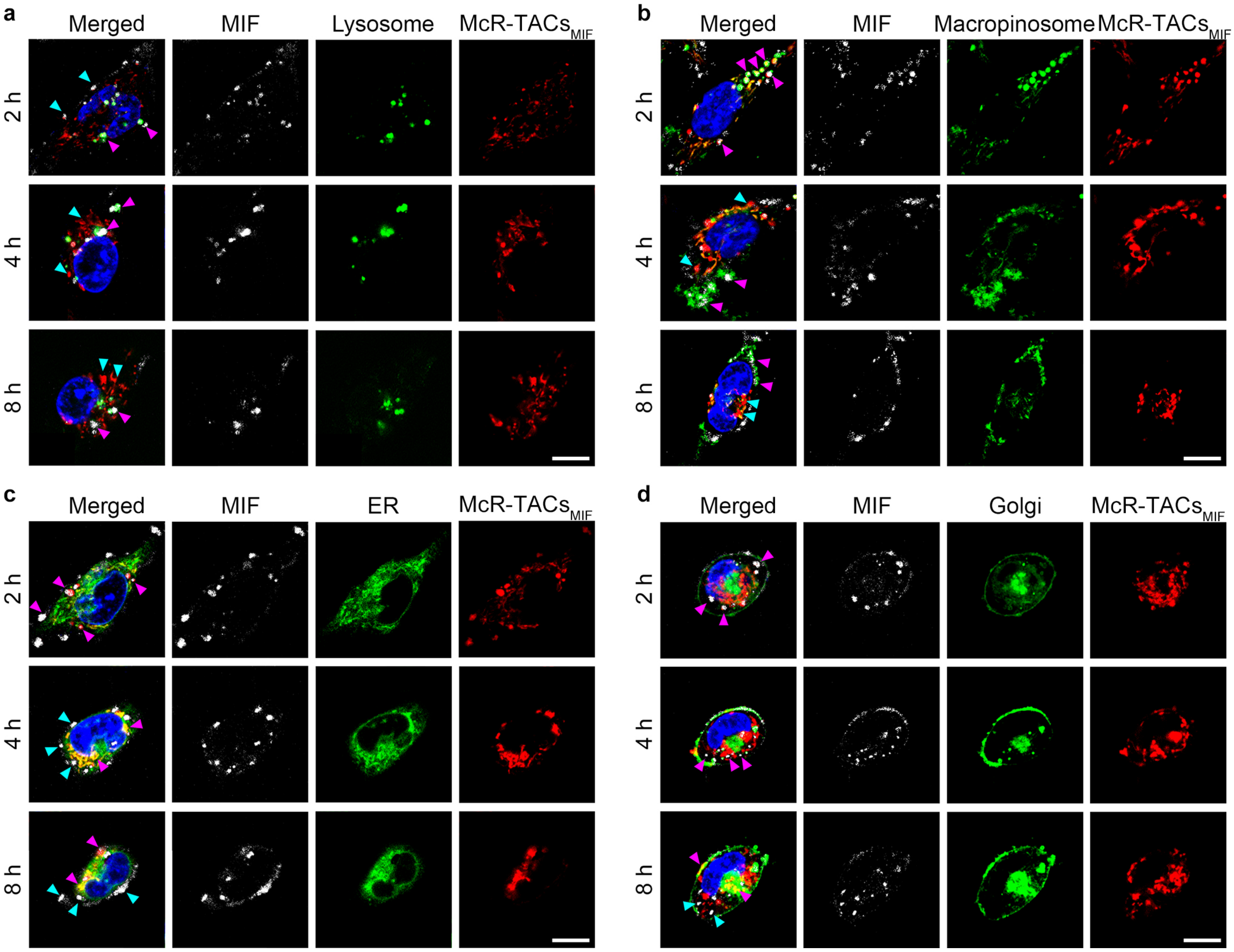


**Fig. S17**. Confocal imaging of the localization of sulfo-Cy5-McR-TACs$_{MIF}$, Fluorescein-MIF and organelles including lysosomes labeled with lysotracker red (**a**), macropinosomes labeled with TRITC-dextran (**b**), ER labeled with ER tracker red (**c**) and Golgi labeled with TMR ceramide (**d**) at indicated times. The purple arrowheads point to the co-localized signals and the blue arrowheads point to the independent signals. Scale bar, 20 μm.

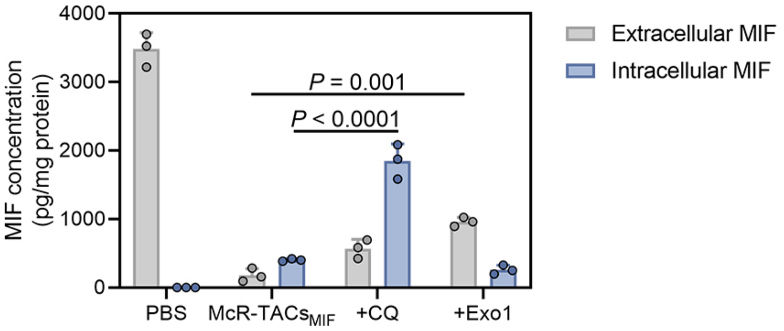


**Fig. S18.** ELISA analysis of the concentrations of both extracellular and intracellular MIF in 4T1 cells treated with McR-TACs$_{MIF}$ with or without indicated inhibitors (*n* = 3 per group). Data are presented as the mean values ± s.d. and were analyzed by a two-way ANOVA, followed by Tukey's multiple comparisons test.

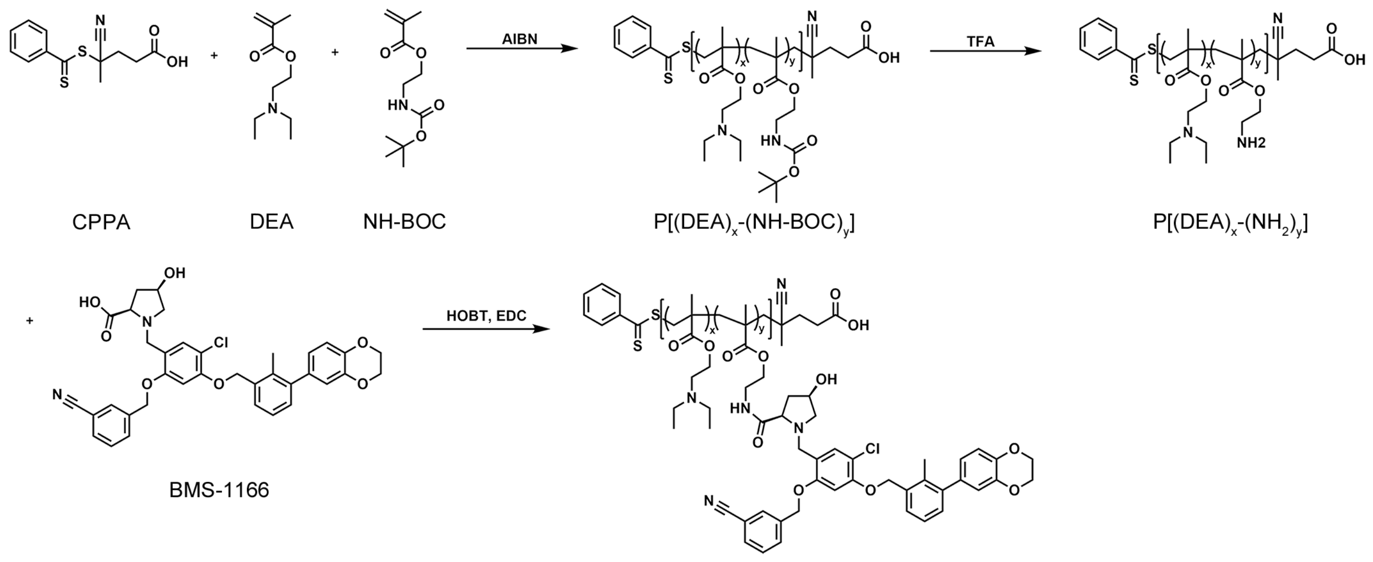


**Fig. S19**. Synthetic routes of N-TACs$_{PD-L1}$.

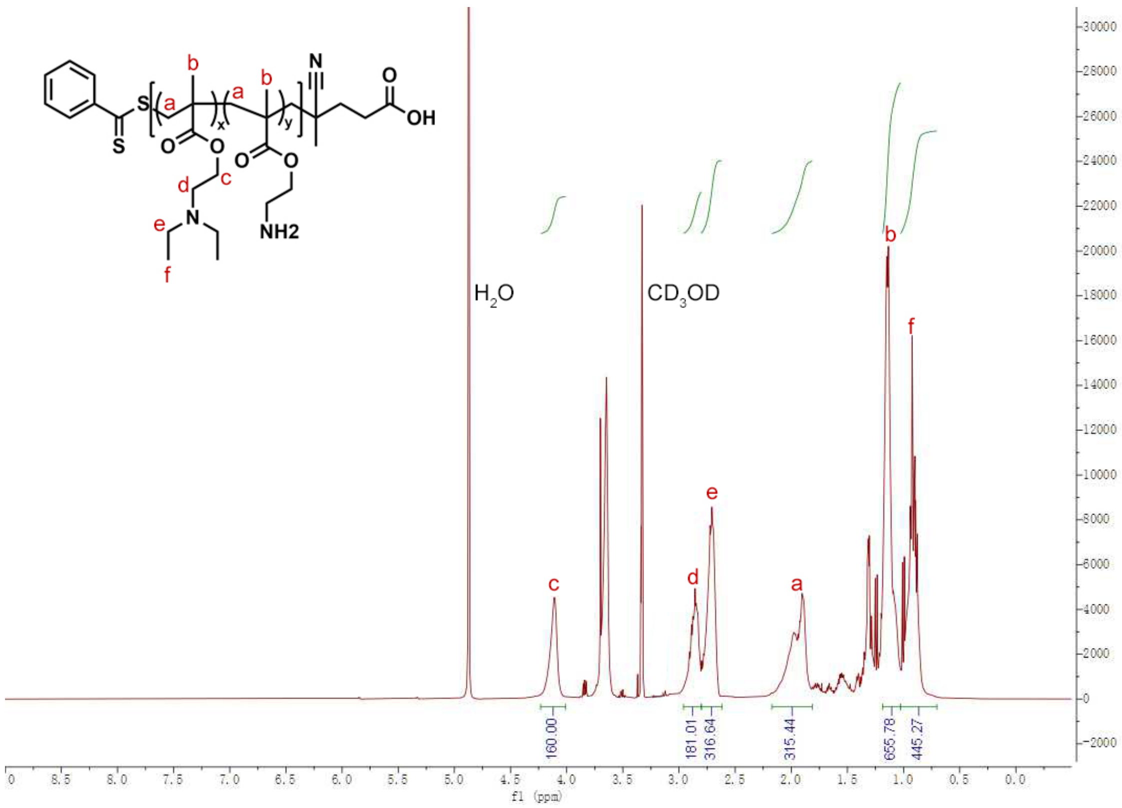


**Fig. S20**. [1]H-NMR spectrum of P[$(DEA)_x$-$(NH_2)_y$] in $CD_3OD$.

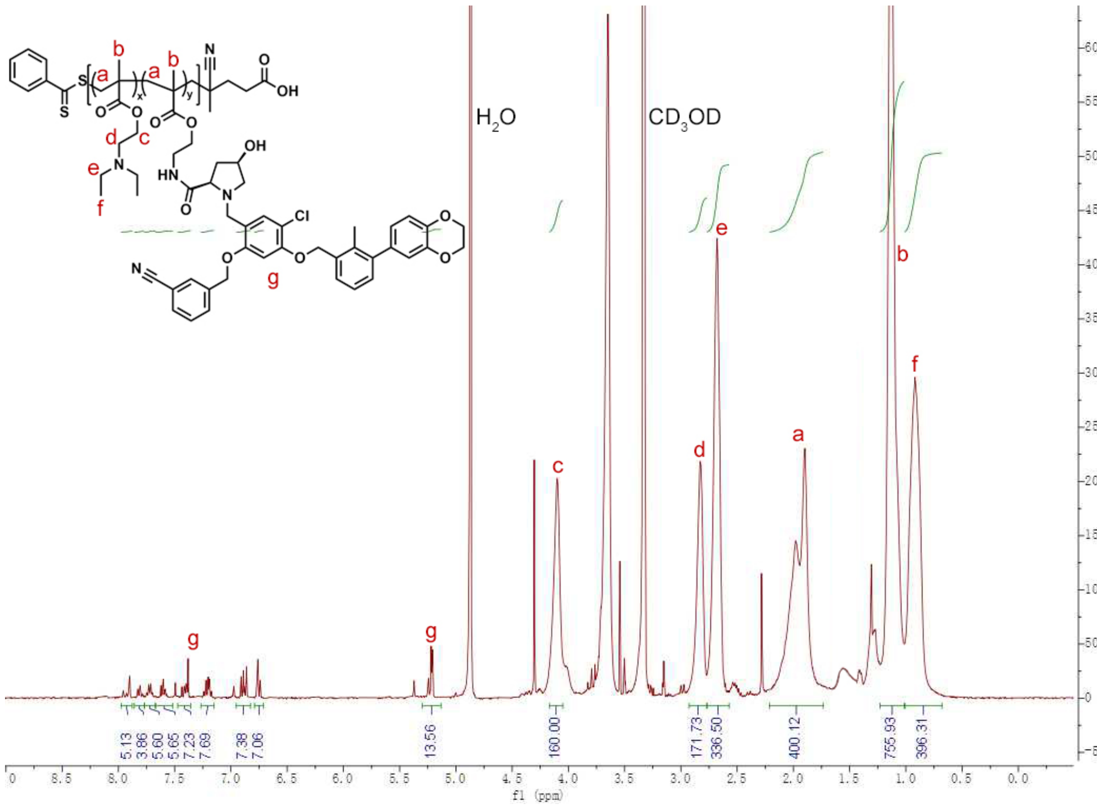


**Fig. S21**. [1]H-NMR spectrum of N-TACs$_{PD-L1}$ in $CD_3OD$.

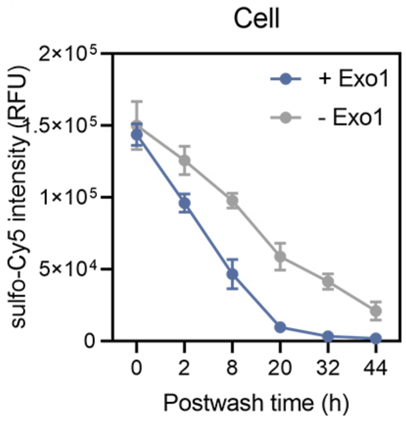


**Fig. S22**. Quantified fluorescent intensity of sulfo-Cy5-labeled McR-TACs in MDA-MB-231 cells at indicated times after washing ($n$ = 3). Data are presented as the mean values ± s.d.

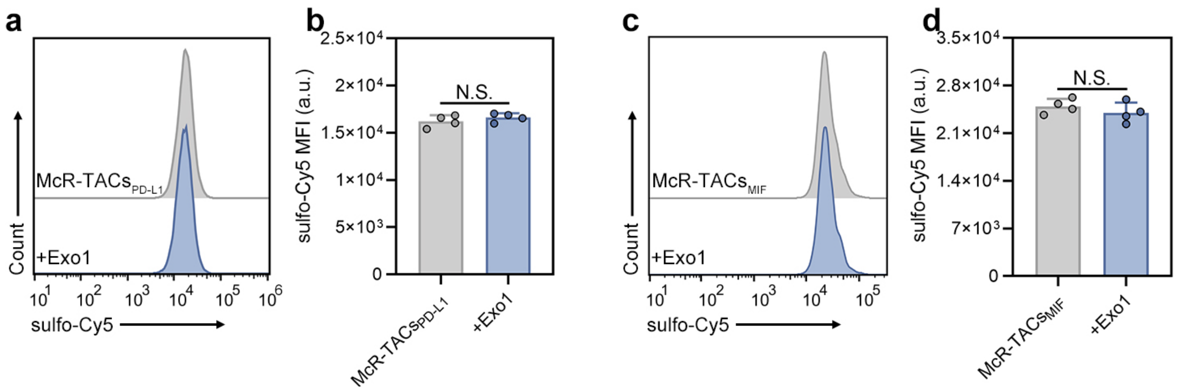


**Fig. S23. The influence of Exo1 on the uptake of McR-TACs.** Flow cytometry analysis of the internalizations of sulfo-Cy5-McR-TACs$_{PD-L1}$ (**a**, **b**) and sulfo-Cy5-MIF (**c**, **d**) in MDA-MB-231 cells (*n* = 4 per group). Data are presented as the mean values ± s.d. and were analyzed by two-tailed unpaired t-test. Not significant (N.S.).

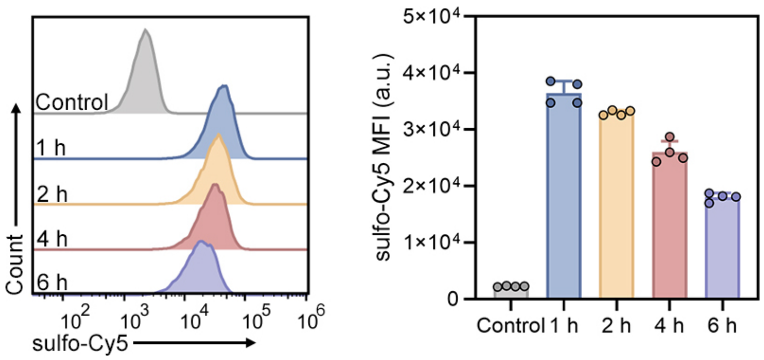


**Fig. S24**. Flow cytometry analysis of sulfo-Cy5-MIF degradation in MDA-MB-231 cells at indicated times after washout ($n$ = 4 per group). Data are presented as the mean values ± s.d.

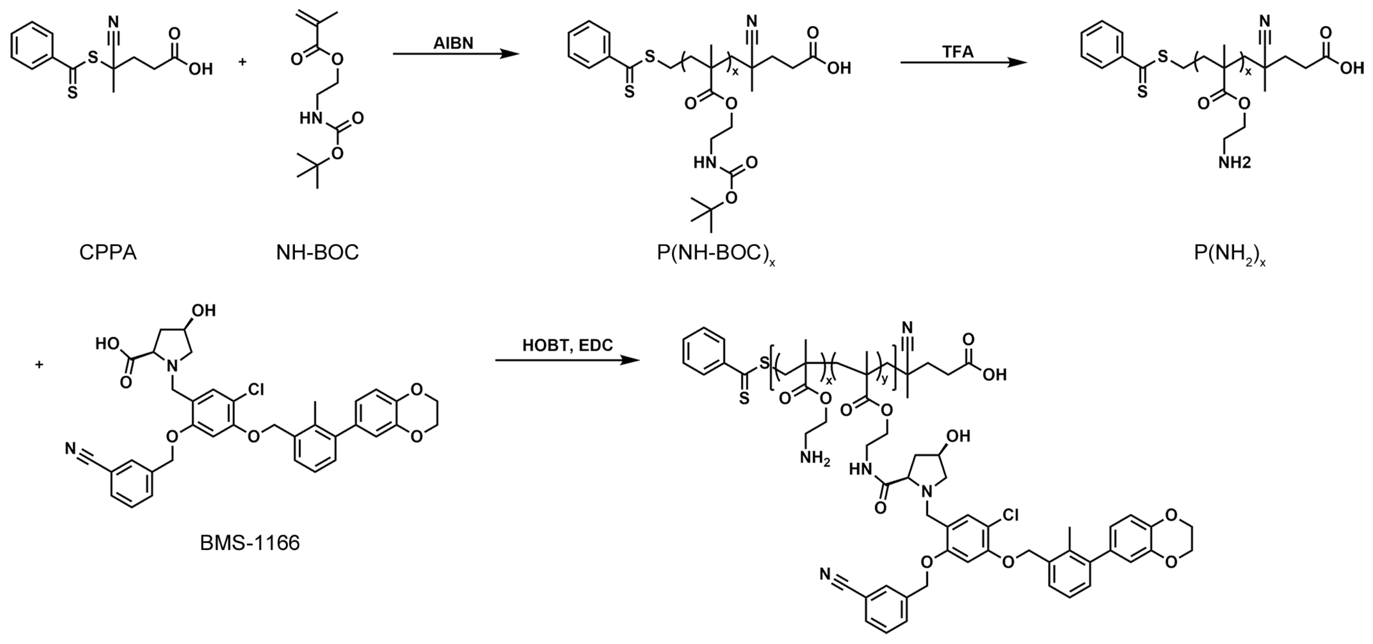


**Fig. S25**. Synthetic routes of AnR-TACs$_{PD-L1}$.

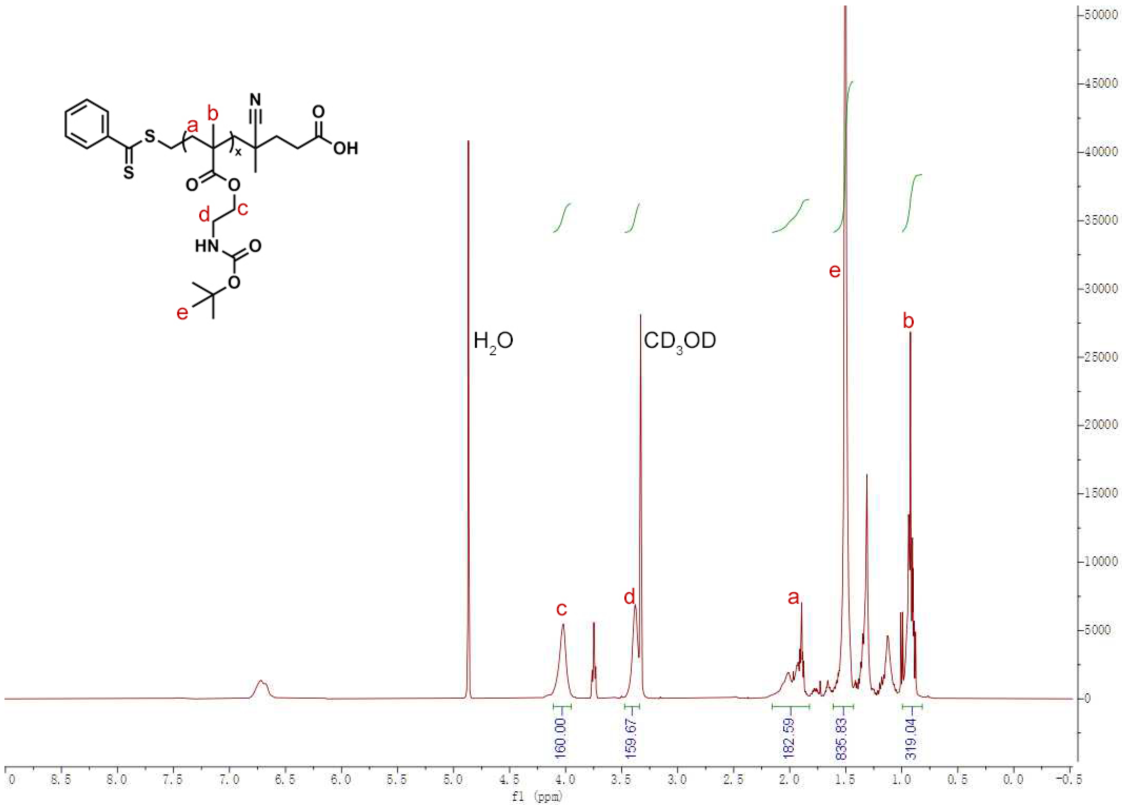


**Fig. S26**. $^{1}$H-NMR spectrum of $P(NH\text{-}BOC)_x$ in $CD_3OD$.

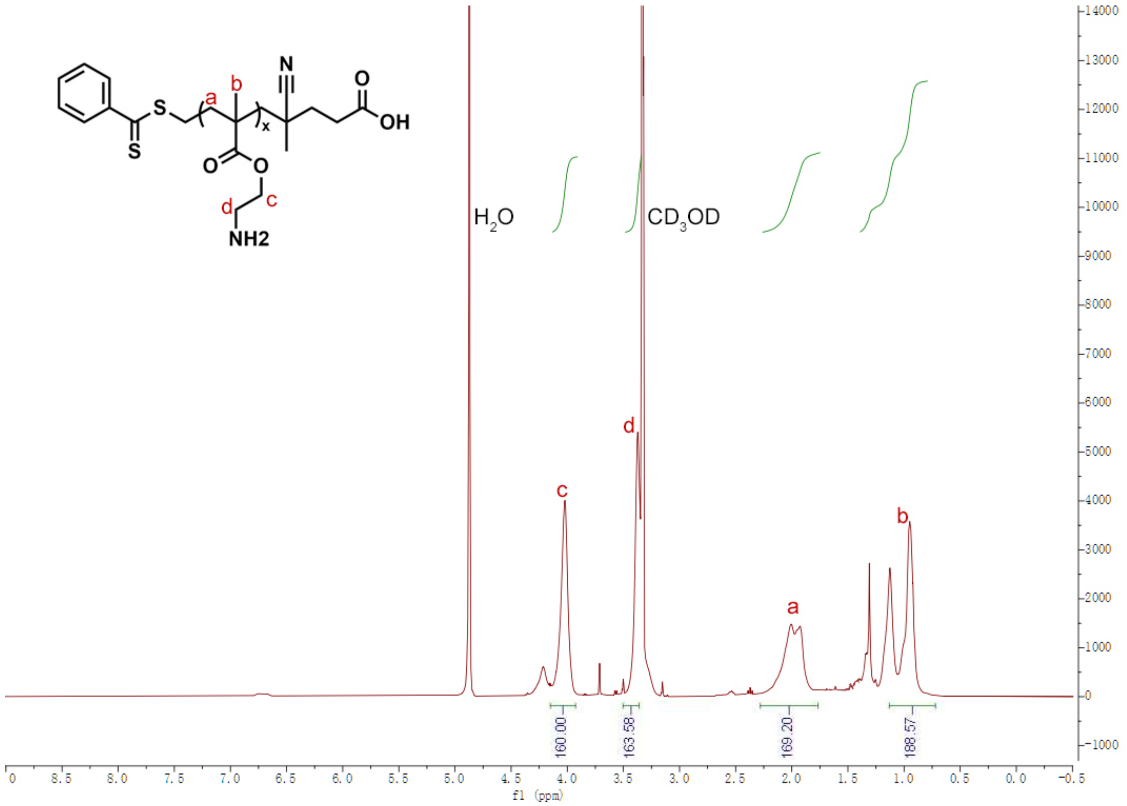


**Fig. S27**. [1]H-NMR spectrum of $P(NH_2)_x$ in $CD_3OD$.

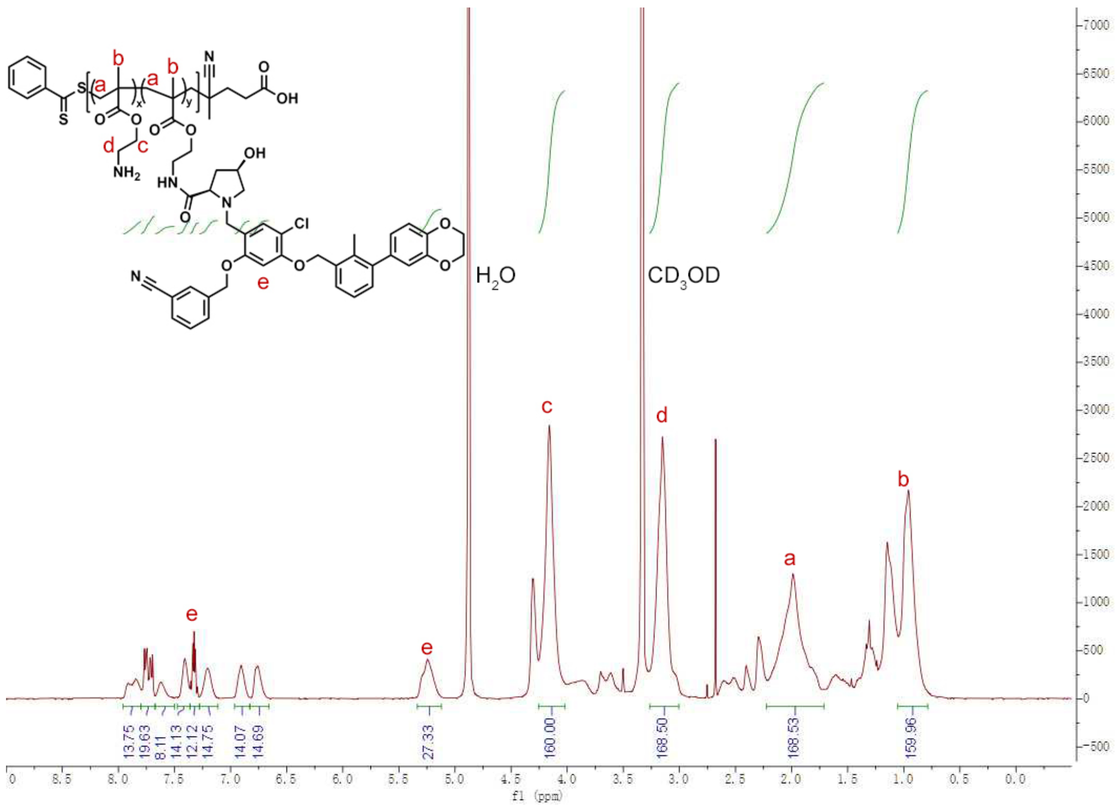


**Fig. S28**. [1]H-NMR spectrum of AnR-TACs$_{PD-L1}$ in $CD_3OD$.

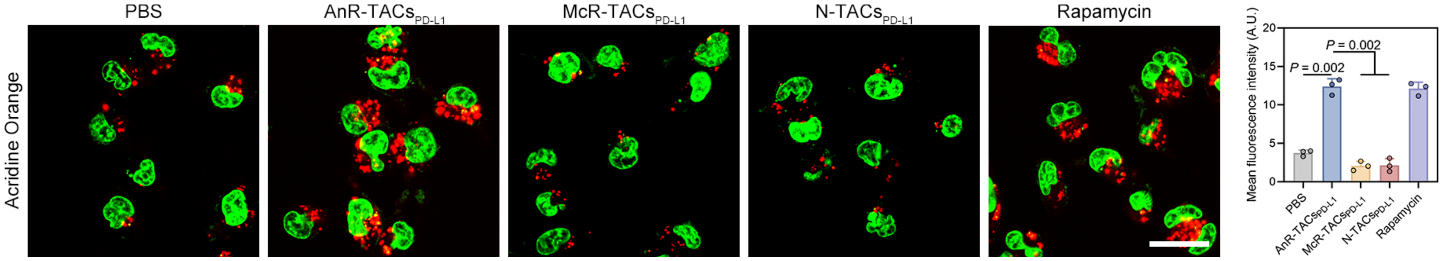


**Fig. S29**. Confocal imaging and semi-quantification of autophagosomes in MDA-MB-231 cells treated with PBS, AnR-TACs$_{PD-L1}$, N-TACs$_{PD-L1}$, McR-TACs$_{PD-L1}$ and rapamycin for 24 h ($n$ = 3 per group). Scale bar, 40 µm. Data are presented as the mean values ± s.d. and were analyzed by a one-way ANOVA, followed by a Tukey's multiple comparisons test.

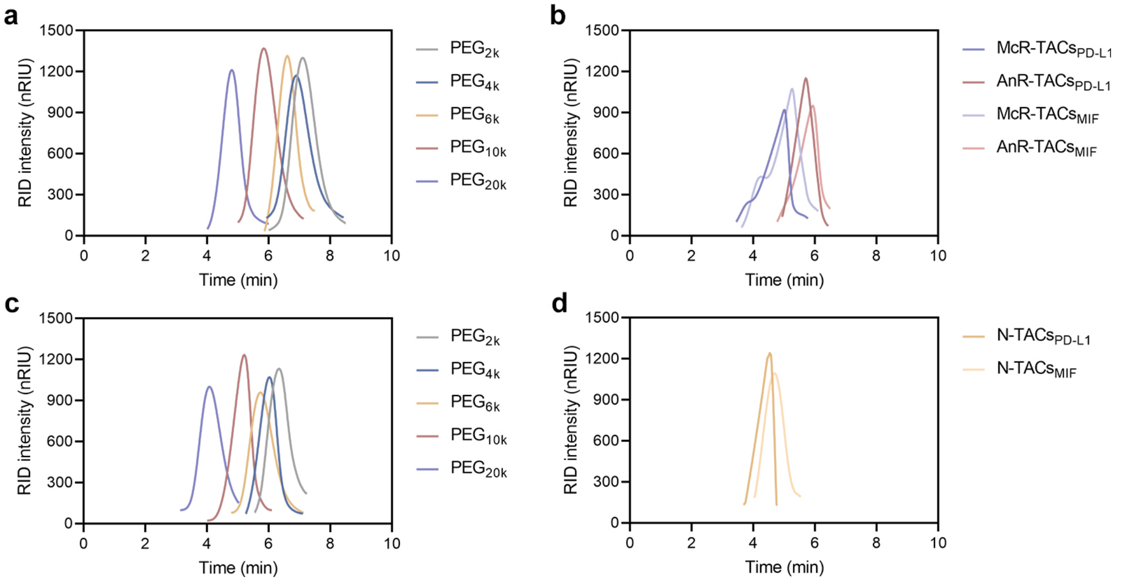


**Fig. S30**. The GPC traces of standard polyethylene glycol (**a**, **c**) and different groups of LYTACs (**b**, **d**) in aqueous (**a**, **b**) and organic (**c**, **d**) phases.

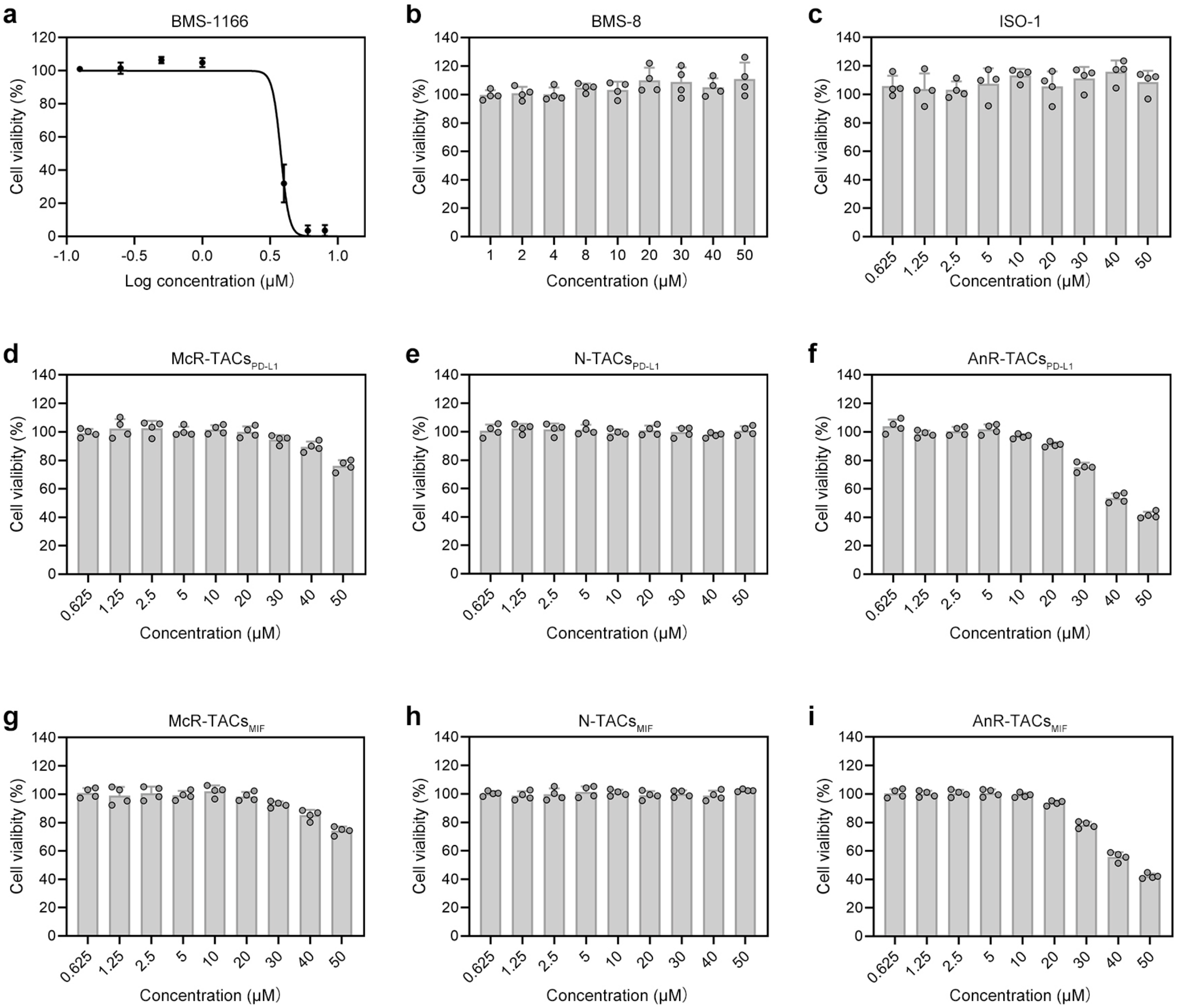


**Fig. S31**. In vitro cytotoxicity of warheads including BMS-1166 (**a**), BMS-8 (**b**) and ISO-1 (**c**) and different groups of LYTACs including McR-TACs$_{PD-L1}$ (**d**), N-TACs$_{PD-L1}$ (**e**), AnR-TACs$_{PD-L1}$ (**f**), McR-TACs$_{MIF}$ (**g**), N-TACs$_{MIF}$ (**h**) and AnR-TACs$_{MIF}$ (**i**) on MDA-MB-231 cells after 24 h treatment ($n$ = 4 per group). Data are presented as the mean values ± s.d.

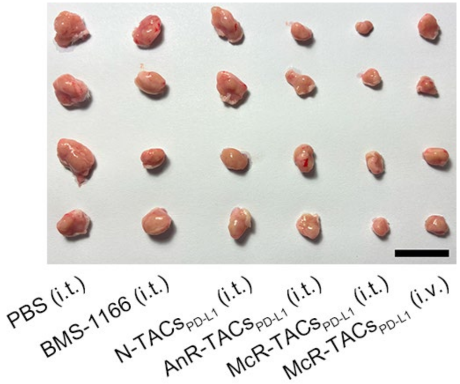


**Fig. S32**. Images of tumors from TNBC-bearing mice with orthotopic 4T1-hPD-L1 breast cancer in different treatment groups ($n$ = 4 per group). Scale bar, 2 cm.

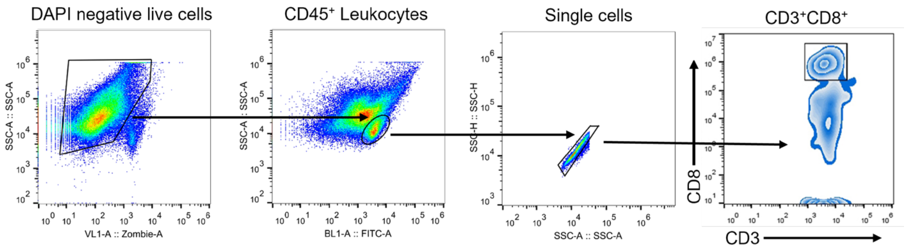


**Fig. S33**. Gating strategy for flow cytometry analysis of $CD45^+$ $CD3^+$ $CD8^+$ T cells in tumor tissues ($n$ = 4 mice).

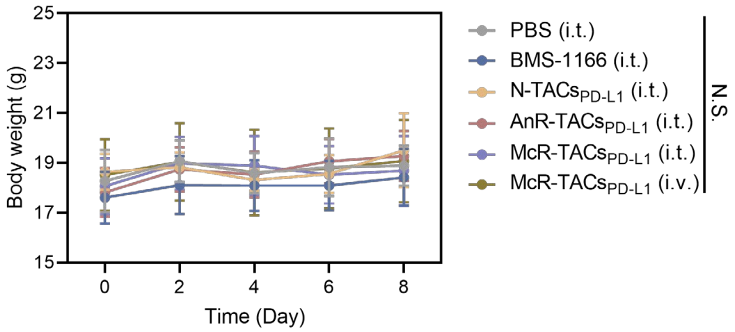


**Fig. S34**. Body weight measurements every two days during treatments ($n$ = 8 per group). Data are presented as the mean values ± s.d.. and were analyzed by a two-way ANOVA, followed by a Tukey's multiple comparisons test. Not significant (N.S.).

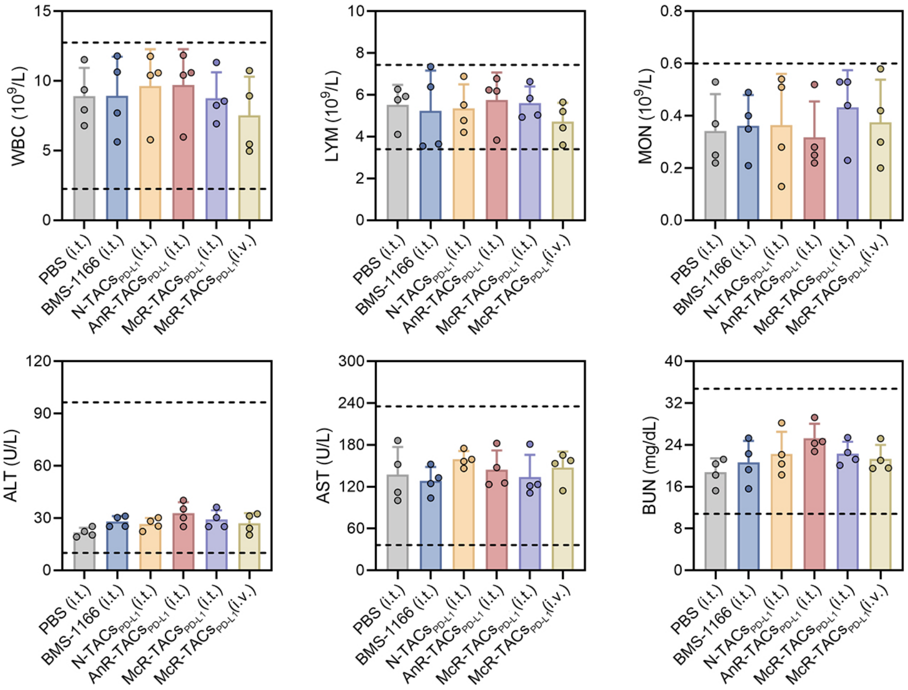


**Fig. S35**. Hematological analysis of TNBC-bearing mice after treatments for PD-L1 degradation. The dashed lines represent the upper and lower limits of normal values ($n$ = 4 per group). Data are presented as the mean values ± s.d.

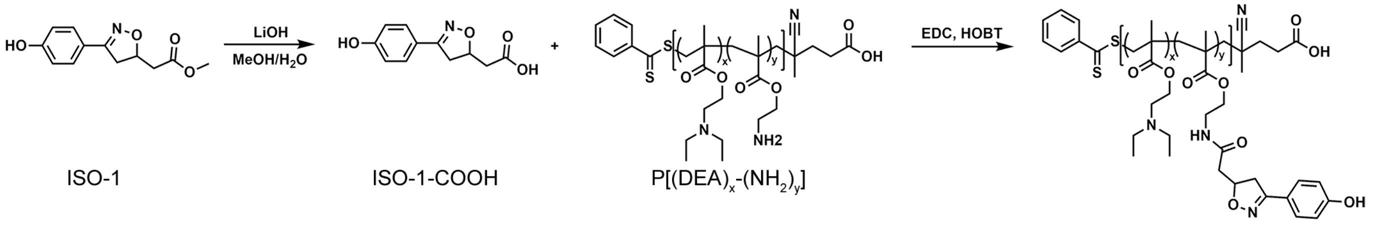


**Fig. S36**. Synthetic routes of N-TACs$_{MIF}$.

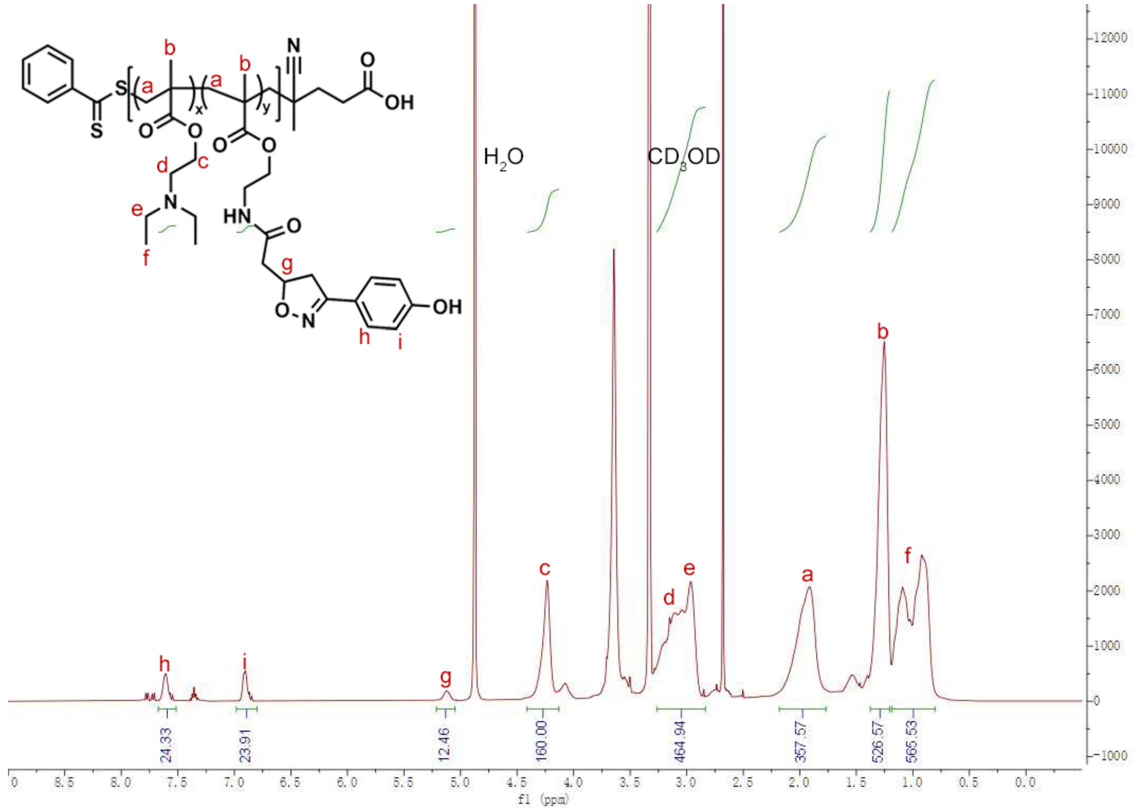


**Fig. S37**. $^{1}$H-NMR spectrum of N-TACs$_{MIF}$ in $CD_3OD$.

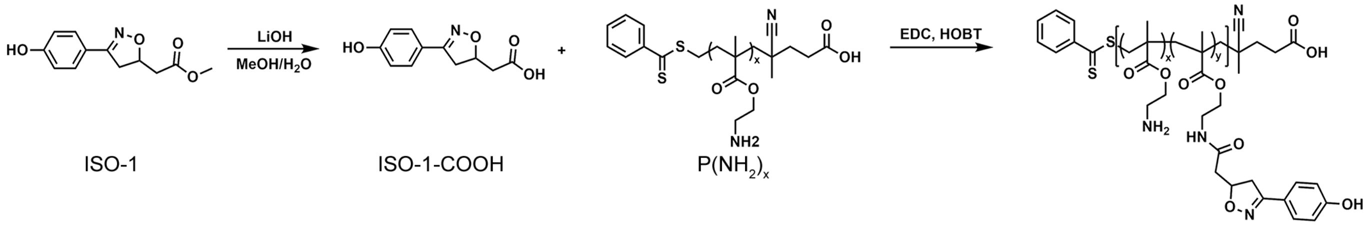


**Fig. S38**. Synthetic routes of AnR-TACs$_{MIF}$.

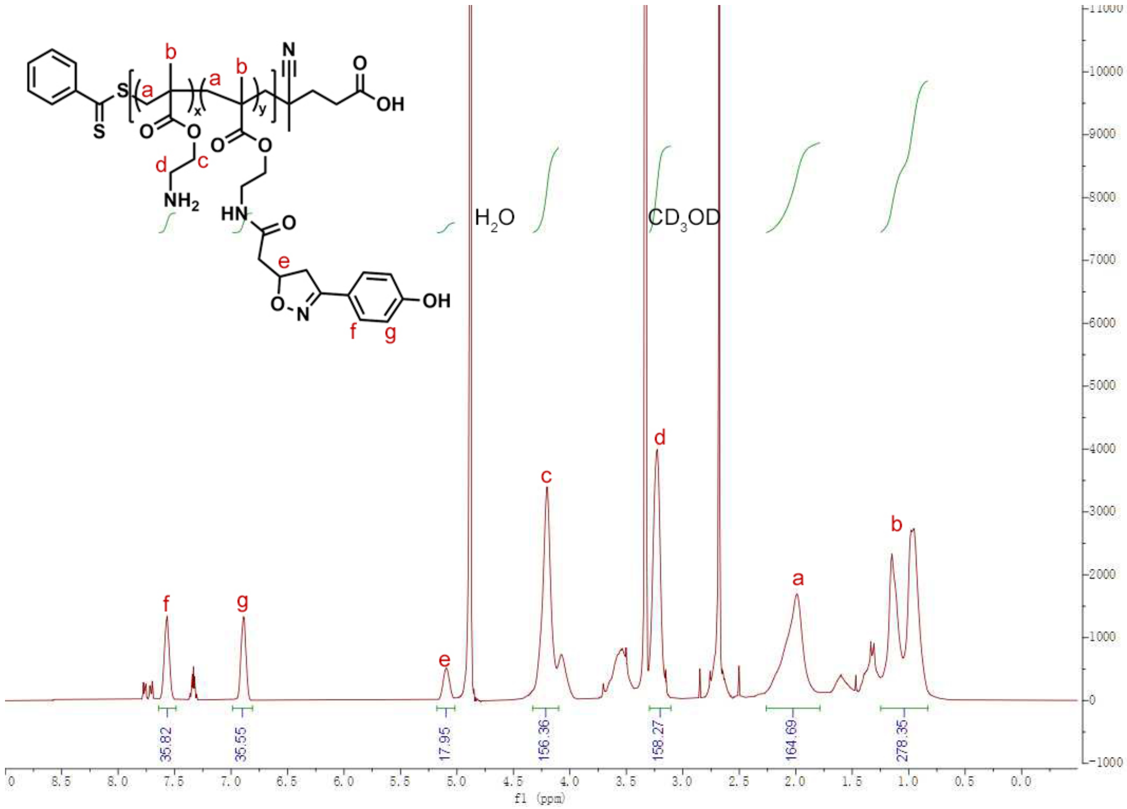


**Fig. S39**. $^{1}$H-NMR spectrum of N-TACs$_{MIF}$ in $CD_3OD$.

### 3. Supplementary tables

**Table. S1**. The Mn and PDI of different LYTACs calculated by GPC.

| Polymer | Mn | PDI |
|---|---|---|
| McR-TACs$_{PD-L1}$ | 19956 | 1.48 |
| McR-TACs$_{MIF}$ | 16904 | 1.42 |
| AnR-TACs$_{PD-L1}$ | 12898 | 1.37 |
| AnR-TACs$_{MIF}$ | 11216 | 1.35 |
| N-TACs$_{PD-L1}$ | 17231 | 1.27 |
| N-TACs$_{MIF}$ | 15149 | 1.29 |

**Table. S2**. The estimated number of monomers in different LYTACs.

| Polymer | x | y |
|---|---|---|
| McR-TACs$_{PD\text{-}L1}$ | 74 | 6 |
| McR-TACs$_{MIF}$ | 68 | 10 |
| AnR-TACs$_{PD\text{-}L1}$ | 71 | 5 |
| AnR-TACs$_{MIF}$ | 61 | 11 |
| N-TACs$_{PD\text{-}L1}$ | 80 | 4 |
| N-TACs$_{MIF}$ | 65 | 10 |

**Table. S3**. The sequence of siRNA used in this study.

| Target gene symbol | Sequence (5'→3') | |
|---|---|---|
| RAC1 | Sense | GCCUUCUUAAAGCCUUAUUtt |
| | Antisense | AAUAAGGCUUUAAGAAGGCtt |
| KRAS | Sense | GGUGACUUAGGUUCUAGAUtt |
| | Antisense | AUCUAGAACCUAAGUCACCtt |
